# Combinatorial expression of gamma-protocadherins regulates synaptic connectivity in the mouse neocortex

**DOI:** 10.1101/2021.11.03.467093

**Authors:** Yi-Jun Zhu, Cai-Yun Deng, Liu Fan, Ya-Qian Wang, Hui Zhou, Hua-Tai Xu

**Affiliations:** Institute of Neuroscience and State Key Laboratory of Neuroscience, CAS Center for Excellence in Brain Science and Intelligence Technology, Chinese Academy of Sciences, Shanghai, 200031, China; Lingang Laboratory, Shanghai Center for Brain Science and Brain-Inspired Intelligence Technology, Shanghai 201210, China; University of Chinese Academy of Sciences, Beijing 100049, China

**Author notes:** Correspondence should be addressed to H. -T. X.

## Abstract

In the process of synaptic formation, neurons must not only adhere to specific principles when selecting synaptic partners but also possess mechanisms to avoid undesirable connections. Yet, the strategies employed to prevent unwarranted associations have remained largely unknown. In our study, we have identified the pivotal role of combinatorial clustered protocadherin gamma (γ-PCDH) expression in orchestrating synaptic connectivity in the mouse neocortex. Through 5-prime end single-cell sequencing, we unveiled the intricate combinatorial expression patterns of γ-PCDH variable isoforms within neocortical neurons. Furthermore, our whole-cell patch-clamp recordings demonstrated that as the similarity in this combinatorial pattern among neurons increased, their synaptic connectivity decreased. Our findings elucidate a sophisticated molecular mechanism governing the construction of neural networks in the mouse neocortex.

## Introduction

The precision of synaptic connections is vital for the functioning of neural circuits(Yogev and Shen, 2014). Cell adhesion molecules play crucial roles in the specificity of synapse formation(Duan et al., 2014, Serizawa et al., 2006, Tan et al., 2015, Rawson et al., 2017, Sytnyk et al., 2017, Berns et al., 2018, Jang et al., 2017, Courgeon and Desplan, 2019). However, how to achieve such specificity at the microcircuit level remains an open question. The unique expression pattern of clustered protocadherins (cPCDH) leads to millions of possible combinations of cPCDH isoforms on the neuron surface(Kaneko et al., 2006, Esumi et al., 2005), effectively serving as a distinctive barcode for each neuron(Yagi, 2012). Notably, the absence of γ-PCDH does not induce general abnormalities in the development of the cerebral cortex, including cell differentiation, migration, and survival(Wang et al., 2002, Garrett et al., 2012). However, γ-PCDH presence has been detected at synaptic contacts(Fernandez-Monreal et al., 2009, Phillips et al., 2003, LaMassa et al., 2021), and its absence has substantial effects on neuronal connectivity(Tarusawa et al., 2016, Kostadinov and Sanes, 2015, Lv et al., 2022). While the homophilic properties of γ-PCDH promote dendritic complexity(Molumby et al., 2016), emerging evidence suggests that it might hinder synapse formation. Previous studies indicate that homophilic interactions, facilitated by large overlapping patterns of cPCDH isoforms on opposing cell surfaces, may lead to intercellular repulsion(Rubinstein et al., 2015, Brasch et al., 2019, Honig and Shapiro, 2020, Lefebvre et al., 2012). Consistent with the repulsion concept, the absence of γ-PCDH results in significantly more dendritic spines and inhibitory synapse densities in neocortical neurons(Molumby et al., 2017, Steffen et al., 2021). Paralleling this, neurons overexpressing one of the γ-PCDH isoforms exhibit significantly fewer dendritic spines(Molumby et al., 2017). Furthermore, the absence of the clustered PCDH augments local reciprocal neural connection between lineage-related neurons in the neocortex(Tarusawa et al., 2016, Lv et al., 2022), even when sister cells exhibit more similar expression patterns of γ-PCDH isoforms(Lv et al., 2022).

Intriguingly, while γ-PCDH appears to have “contradictory” effects on dendritic complexity and dendritic spines, it negatively influences synapse formation in the forebrain. It’s important to note that each neuron expresses multiple isoforms of γ-PCDH(Kaneko et al., 2006, Lv et al., 2022). What is the impact of this combinatorial expression on synapse formation? In this study, using 5-prime end (5’-end) single-cell sequencing, we revealed the diversified combinatorial expression of γ-PCDH isoforms in neocortical neurons. Through multiple whole-cell patch-clamp recordings after the sequential *in utero* electroporation, we discovered that the combinatorial expression of γ-PCDHs empowers neurons to decide which partners to refrain from forming synapses with, rather than merely determining which ones to engage in synaptogenesis with.

## Results

### The diversified combinatorial expression pattern of γ-PCDHs in neocortical neurons revealed by 5’-end single-cell sequencing

The gamma isoform of cPCDHs (γ-PCDHs) is critical for synaptic connectivity(Kostadinov and Sanes, 2015, Tarusawa et al., 2016, Lv et al., 2022). To determine the role of γ-PCDH in the neocortex, we examined their expression in the neocortical neurons of postnatal mice. Existing research has suggested that clustered protocadherins are expressed stochastically in Purkinje cells and olfactory sensory neurons(Hirayama et al., 2012, Toyoda et al., 2014, Mountoufaris et al., 2017). In this study, we harnessed the power of 5’-end single-cell RNA sequencing to precisely identify γ-PCDH isoforms by focusing on their variable exon, exon 1, where they differ from each other(Kohmura et al., 1998, Wu and Maniatis, 1999). Given that the second postnatal week is the critical stage for synapse formation in the rodent neocortex(Lendvai et al., 2000, Holtmaat and Svoboda, 2009), we chose postnatal day 11 (P11) as the time point for our examination. Following reverse transcription and cDNA amplification (Fig. 1-S1A, B), we divided the cDNA into two segments: one designed for the specific amplification of *Pcdhg* mRNAs and the other for the construction of a 5’ gene expression library (Fig. 1-S1C). After the cluster analysis (Fig. 1-S2-4), we collected 6505 neurons from an initial pool of 17438 cells (Fig. 1A and Fig. 1-S1D). For in-depth analysis, we focused on neurons expressing more than 10 unique molecular identifiers (UMI) for all γ-PCDH isoforms (cutoff >1 for each type of individual isoform) (Fig. 1B, C). We observed the near-ubiquitous expression of “C-type” isoforms, specifically C3, C4, and C5 (Fig. 1D). It’s important to note that the fraction of cells expressing “C-type” isoforms was significantly higher when compared to “variable” isoforms (Fig. 1D and Fig. 1-S1E), which is consistent with findings from a previous study(Toyoda et al., 2014).

**Figure 1:**
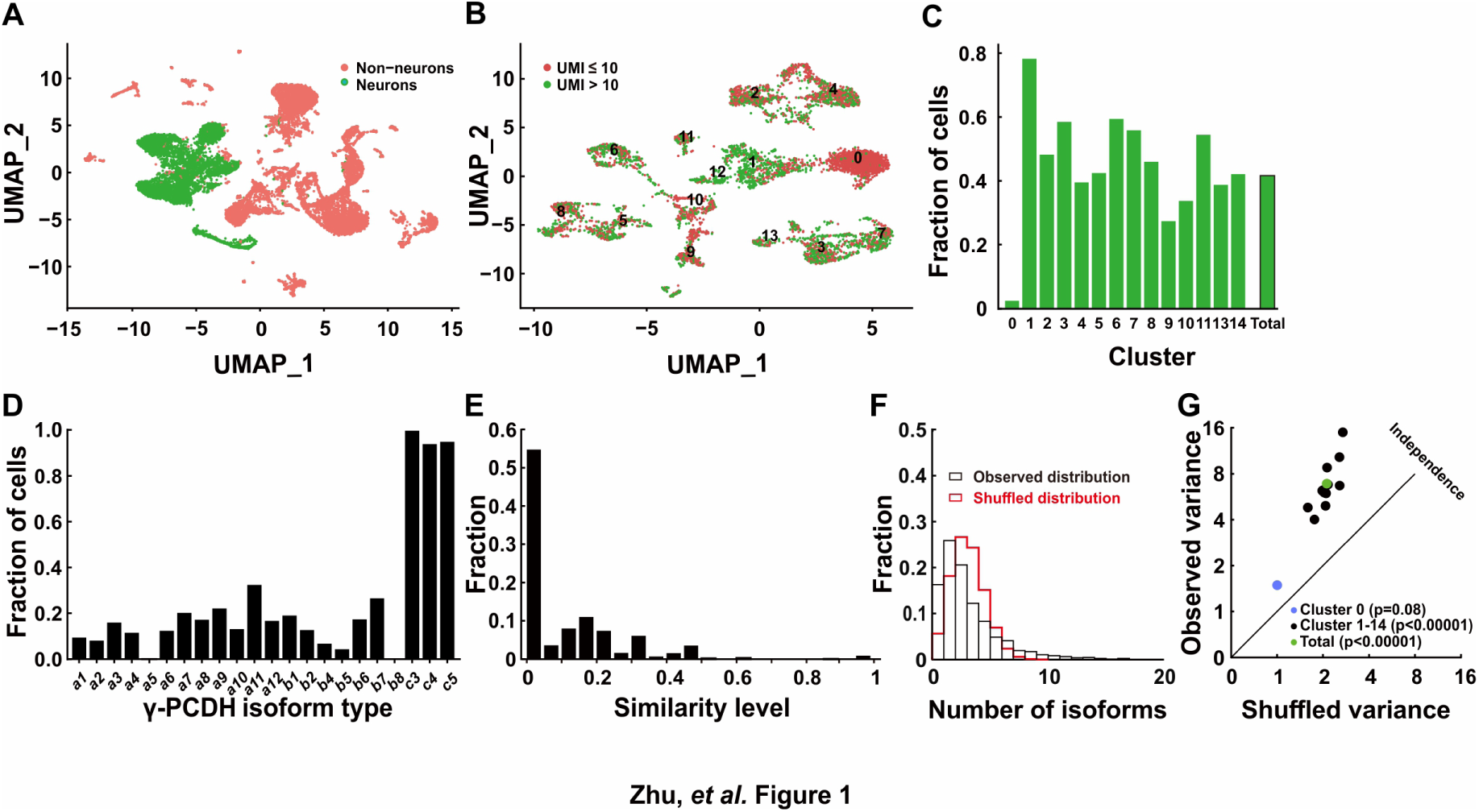
Diversified expression of *Pcdhg* isoforms in neocortical neurons. (**A**) UMAP analysis displaying 17,438 cells obtained through 5’-end single-cell sequencing after data cleanup and doublet removal. Neurons are depicted as green dots, while non-neuronal cells are marked in red. (**B**) UMAP clusters of all neurons categorized by the UMI cutoff. Red dots denote cells with fewer than 10 total UMIs of *Pcdhg* (n=3,671), and green dots denote cells with more than 10 UMIs (n=2,834). (**C**) Fractions of neurons with more than 10 UMIs in different clusters. (**D**) Distribution of neurons expressing different *Pcdhg* isoforms in the neocortex. (**E**) Fraction distribution illustrating similarity levels in the combinatorial expression of *Pcdhg* variable isoforms among neurons. Similarity levels were calculated as ^𝐴𝐴∩𝐵𝐵^. (**F**) Observed distribution (black) of the fraction of cells expressing varying numbers of isoforms across all neurons. The red curve represents the shuffled distribution generated under the hypothesis of stochastic isoform expression. (**G**) The variance difference between the observed and shuffled fraction distribution of cells from all clusters.

We proceeded to conduct a pairwise analysis to assess the similarity of “variable” isoforms among neurons regarding the single-cell expression pattern of γ-PCDHs. This extensive analysis revealed that the majority of neocortical neurons from all clusters exhibited very low similarity level (Fig. 1E, Fig. 1-S5 and 1-S6). This finding strongly suggests distinct combinatorial expression patterns among these neurons. Given that all our electrophysiological recordings were carried out on pyramidal neurons in the layer 2/3 of the neocortex, we conducted more detailed analysis, including the examination of detailed expression of γ-PCDHs in individual neurons, specifically in the corresponding cluster 7. All these analyses consistently revealed a diverse range of combinatorial expression patterns among neurons in this cluster, a phenomenon in alignment with the general population of neocortical neurons (Fig. 1-S5).

To further characterize this diversity, we employed variance analysis as per Wada et al.(Wada et al., 2018), to examine the distribution of the number of expressed isoforms per cell across all neurons (Fig. 1F and Fig. 1-S7). As per Wada’s definition of ‘co-occurrence’ (Wada et al., 2018), this primarily indicates potential interactions among different isoform expressions at the population level. This analysis uncovered a subtle but significant co-occurrence of γ-PCDH isoform expression in most neurons (Fig. 1G). Notably, we did not detect any discernible differences among clusters, except for cluster 0, which displayed considerably lower expression (Fig. 1C) and no co-occurrence of γ-PCDH isoforms (Fig. 1G and Fig. 1-S7). In summary, the data derived from 5’-end single-cell sequencing underscore the diverse and complex combinatorial expression of γ-PCDHs within the majority of neocortical neurons.

### The absence of γ-PCDHs increases local synaptic connectivity among pyramidal neurons in the neocortex

To delve into the function of γ-PCDH in the synaptic formation among neocortical neurons, we conducted paired recordings on pyramidal neurons in the layer 2/3 of the neocortex from *Pcdhg* conditional knockout (cKO) mice. These genetically engineered mice were created by crossing *Pcdhg ^flox/flox^* mice(Lefebvre et al., 2008, Prasad et al., 2008) with *Nex-cre* mice(Goebbels et al., 2006). This genetic combination specifically removed all variable and C-type γ-PCDH isoforms in pyramidal neurons. Our experimental setup involved multiple whole-cell patch-clamp recordings from cortical slices harvested from P9-32 mice, allowing us to measure the connectivity among nearby pyramidal cells (<200 μm between cell somas) in the layer 2/3 of the neocortex by assessing the presence of evoked monosynaptic responses (Fig. 2 and Fig. 2-S1).

**Figure 2:**
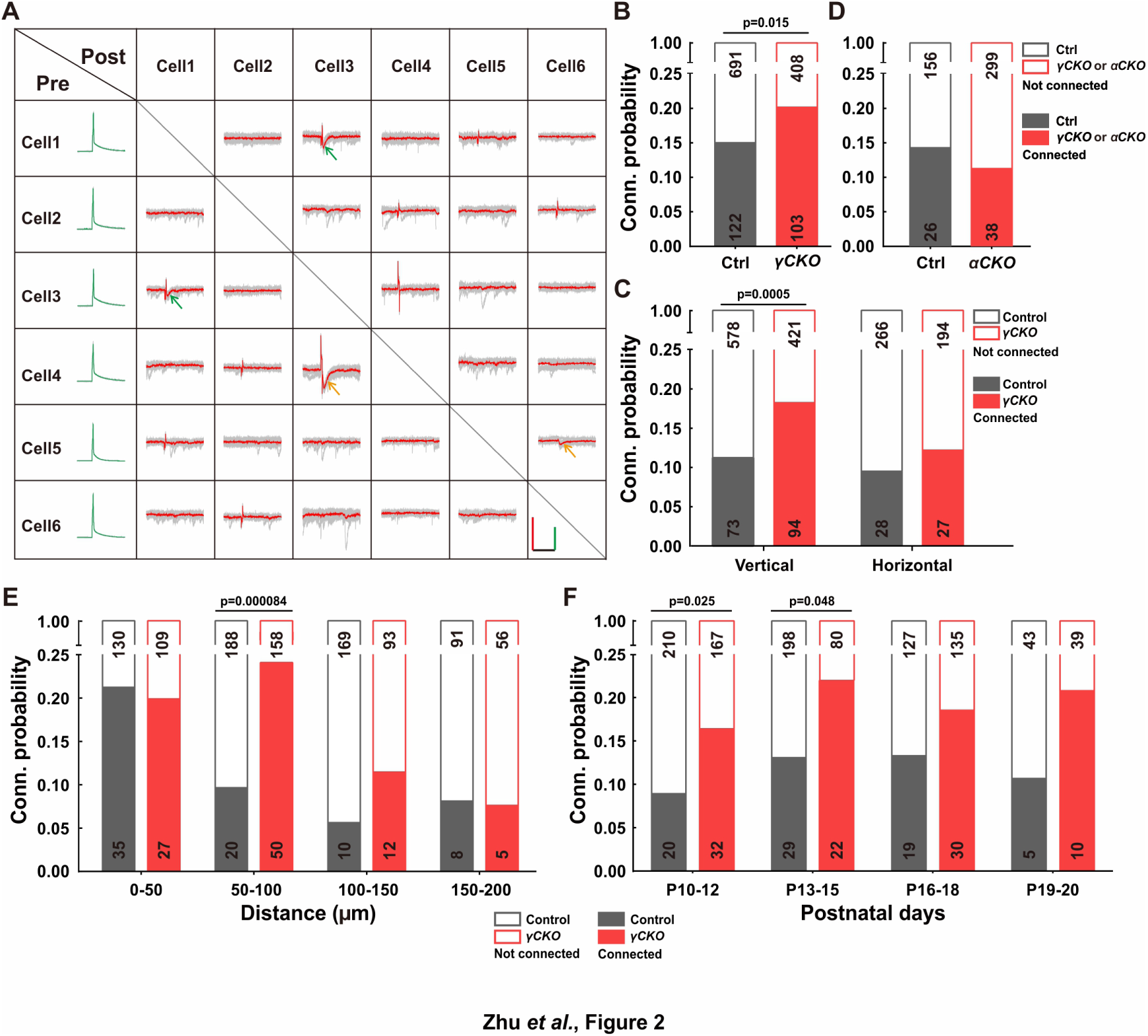
Increased Synaptic Connectivity in the Absence of γ-PCDH. (**A**) Representative traces (red/green) from multiple electrode whole-cell patch-clamp recordings conducted on six neurons in layer 2/3 of the barrel cortex. Average traces are shown in red and green, with ten original traces in gray. Positive evoked postsynaptic responses are indicated by arrows. Orange and green arrows denote unidirectional and bidirectional synaptic connections, respectively. Scale bars: 100 mV (green), 50 pA (red), and 50 ms (black). (**B, D**) Connectivity probability among nearby pyramidal cells in the layer 2/3 of the barrel cortex in *Pcdhg* cKO (**B**), *Pcdha* cKO mice (**D**) and their littermate WT controls. γCKO: *Pcdhg flox/flox*::*nex-cre* mice; αCKO: *Pcdha flox/flox*::*nex-cre* mice; Ctrl: *Pcdhg*+/+::*nex-cre* or *Pcdha*+/+::*nex-cre* mice. (**C**) Connectivity probability among vertically or horizontally aligned neurons in the layer 2/3 of the barrel cortex in *Pcdhg* cKO mice and their littermate WT controls. (**E**) Connectivity probability among vertically aligned pyramidal cells as a function of the distance between recorded pairs in *Pcdhg* cKO mice and WT mice. (**F**) Developmental profiling of connectivity probability among vertically aligned neurons in *Pcdhg* cKO mice and WT mice. Chi-square tests were used in **B**-**F** to calculate statistical differences.

In the sample traces featured in the connectivity matrix obtained from six recorded neurons (Fig. 2A and Fig. 2-S1), we observed two neuronal pairs (neuronal pairs 4→3 and 5→6) exhibiting unidirectional monosynaptic connections (indicated by orange arrows), and one pair (neuronal pair 1↔3) displaying bidirectional connections (indicated by green arrows) out of 15 possible pairings. Overall, our analysis revealed that the percentage of connected pairs was notably higher in *Pcdhg* cKO mice (20.2%, 103/511) than in wild-type (WT) mice (15.0%, 122/813) (Fig. 2B). In light of the distinct roles that vertical vs. horizontal axes might play in synaptic organization within the neocortex(Douglas and Martin, 2004), we conducted an additional set of recordings on P10-20 mice, segregating neuron pairs along these axes. In *Pcdhg* cKO mice, we observed a more significant difference in connectivity for vertically aligned cells (18.3%, 94/515 in *Pcdhg* cKO mice vs. 11.2%, 73/651 in WT mice for vertically aligned neuron pairs; 12.2%, 27/221 in *Pcdhg* cKO mice vs. 9.5%, 28/294 in WT mice for horizontally aligned neuron pairs) (Fig. 2C). We also created *Pcdha* cKO mice (Fig. 2-S2-4) and carried out similar experiments focused on vertically aligned neurons. In these mice lacking α-PCDH, we didn’t observe a significant difference (11.3%, 38/337 in *Pcdha* cKO mice vs. 14.3%, 26/182 in WT mice, Fig. 2D).

A more detailed analysis of synaptic connections among vertically aligned neurons in *Pcdhg* cKO mice unveiled that the absence of γ-PCDH expression significantly increased synapse formation between cells separated vertically by 50 to 100 μm (24.0%, 50/208 in *Pcdhg* cKO mice *vs.* 9.6%, 20/208 in WT mice) (Fig. 2E). Notably, this increased synapse formation was detected starting from P10 (16.1%, 32/199 in *Pcdhg* cKO mice *vs.* 8.7%, 20/230 in WT mice for P10-12; 21.6%, 22/102 in *Pcdhg* cKO mice *vs.* 12.8%, 29/227 in WT mice for P13-15). These time points correspond to the period when chemical synapses between cortical neurons become detectable, and this trend persisted throughout our measurements, spanning up to P20 (Fig. 2F). Hence, our findings suggest that γ-PCDHs may play a role in preventing synapse formation from the early stages of neural development.

### Overexpression of γ-PCDHs decreases local synaptic connectivity in the mouse neocortex

To further determine the influence of γ-PCDHs on synaptic connectivity, we overexpressed randomly selected single or multiple γ-PCDH isoforms tagged with fluorescent proteins through *in utero* electroporation in mice. Subsequently, we performed a series of multiple whole-cell patch-clamp recordings to unveil how γ-PCDHs affect synaptic connectivity between neurons situated in the layer 2/3 of the neocortex.

Firstly, to validate the overexpression effect, we conducted a battery of assays, including Real-Time Quantitative Reverse Transcription PCR (qRT-PCR) (Fig. 3A, B), single-cell RT-PCR (Fig. 3C, D), and immunohistochemistry (Fig. 3-S1A to C). The qRT-PCR assays unequivocally confirmed that the electroporated isoforms, as opposed to the non-overexpressed ones from the contralateral side, exhibited significantly higher expression levels (Fig. 3A, B). To assess how many isoforms were typically expressed in a given neuron when multiple plasmids were electroporated, we strategically tagged the first five isoforms with mNeongreen and the sixth with mCherry (Fig. 3-S1A). Employing a probability analysis based on the occurrence of yellow and red-only cells within the total electroporated neuron population (Fig. 3-S1B), we ascertained that each positive neuron expressed an average of 5.6 types of isoforms (Fig. 3-S1C and the top panel of D). This result harmonized with data obtained from single-cell RT-PCR analysis, wherein an average of 5.3 types out of the six electroporated isoforms was detected from 19 neurons (Fig. 3C, D and the bottom panel of Fig. 3-S1D). These single-cell RT-PCR findings further substantiate that the electroporation-introduced isoforms predominate within these neurons (Fig. 3D).

**Figure 3:**
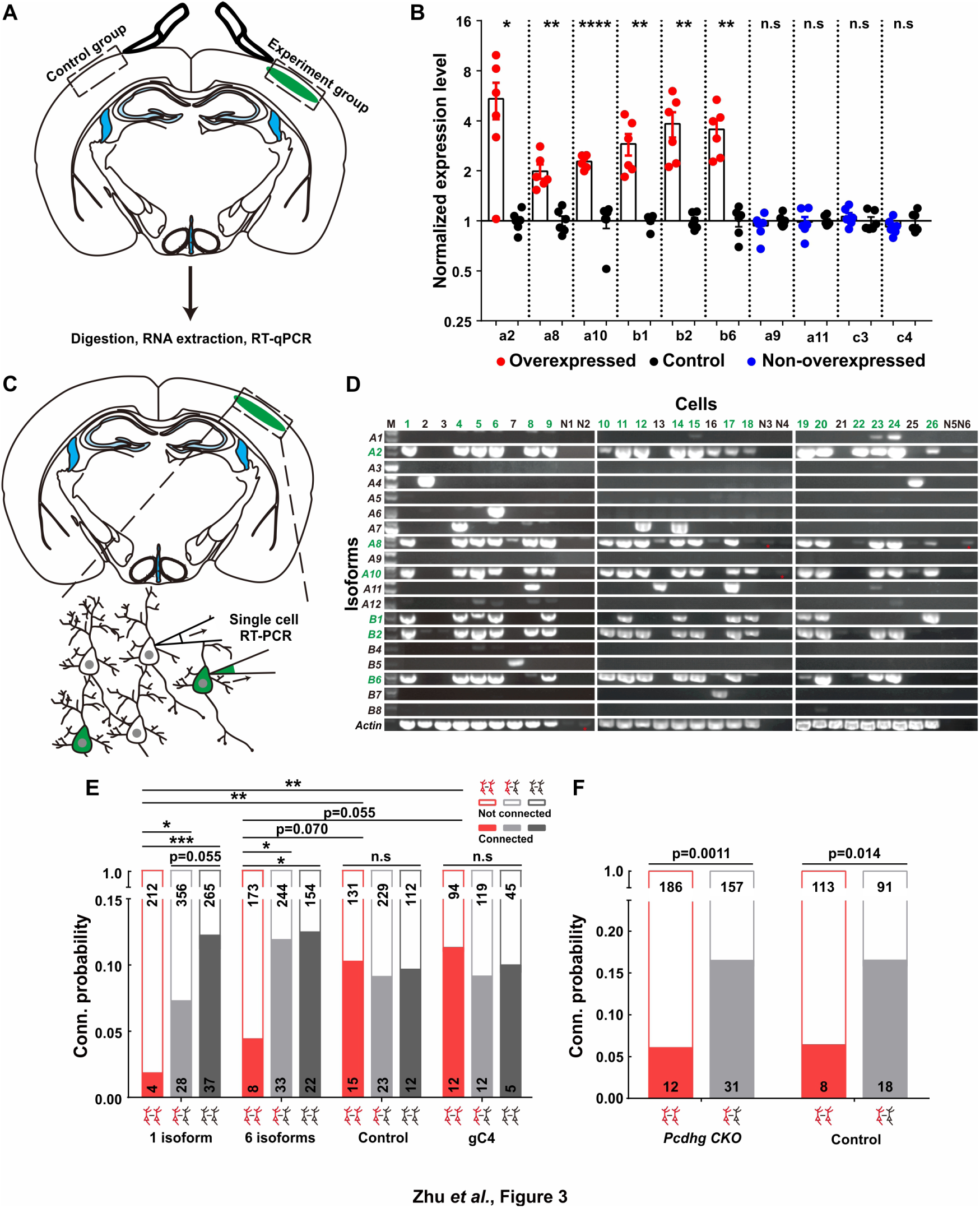
Overexpressing identical variables, but not C4, γ-PCDHs in neurons decreased their synaptic connectivity. (**A**) Schematic illustrating the brain regions selected for qRT-PCR in both experimental and control groups. (**B**) qRT-PCR results showing overexpression levels in electroporated regions. Electroporated isoforms are indicated in red, control isoforms in blue, and contralateral sides used as controls are in black. Statistical analysis was conducted using Student’s t-test, where * indicates p<0.05, ** p<0.01, *** p<0.001, and **** p<0.0001. (**C**) Diagram illustrating the process of cell extraction for single-cell RT-PCR assays. (**D**) Results of single-cell RT-PCR for γ-PCDH isoforms after electroporation. Neurons with fluorescence are highlighted in green, while nearby neurons without fluorescence are in black. Negative controls are labeled as N1-N6. Electroporated isoforms are shown in green, with red stars indicating faint signals in negative controls. (**E**) Impact of overexpressing one or six γ-PCDH isoforms on synaptic connectivity in WT mice. “1 isoform” represents *Pcdhga2*, and “6 isoforms” denote *Pcdhga2*, *Pcdhga8*, *Pcdhga10*, *Pcdhgb1*, *Pcdhgb2*, and *Pcdhgb6*. “gC4” stands for *PcdhgC4*, and “Control” indicates plasmid vector without *Pcdhg* insertion. (**F**) The influence of overexpressing 6 γ-PCDH isoforms on synaptic connectivity in *Pcdhg* cKO mice. The same 6 isoforms used as in (**E**) were employed. *Pcdhg cKO*: *Pcdhg* conditional knockout mice; Control: WT littermates. Statistical differences between groups in (**E**) and (**F**) were determined using the chi-square test and false discovery rate (FDR, Benjamini-Hochberg method) correction.

Since the distribution of neocortical neurons across different layers significantly influences their synaptic connections(He et al., 2015), we meticulously examined the positions of these neurons relative to the pial surface after overexpression. Remarkably, we observed that overexpressing γ-PCDH isoforms did not induce any alterations in cell positioning within the neocortex compared to the control plasmids (Fig. 3-S1E, F). Subsequently, through multiple recordings, we embarked on a quest to assess the impact of γ-PCDHs on synapse formation. Intriguingly, overexpressing one or six γ-PCDH variable isoforms in neurons significantly diminished the rate of synaptic connections among them (10.3%, 15/146 in expressing control plasmids; 1.9%, 4/216 in overexpressing one isoform; and 4.4%, 8/181 in overexpressing six isoforms, red bars in Fig. 3E). However, when overexpressing γ-PCDH C4, we did not observe any significant effect on the synaptic connection rate (11.3%, 12/106 in neurons overexpressing γ-PCDH C4, Fig. 3E).

To further exclude the potential influence of C-type γ-PCDHs in synapse formation, we employed a similar strategy in *Pcdhg* cKO mice, electroporating six variable isoforms. Remarkably, this overexpression also led to a reduction in the connection rate in *Pcdhg* cKO mice (6.1%, 12/198 in overexpressing six isoforms vs. 16.5%, 31/188 in expressing control plasmids), mirroring the outcome observed in WT littermate mice (6.6%, 8/121 in overexpressing six isoforms vs. 16.5%, 18/109 in expressing control plasmids) (Fig. 3F). These compelling observations collectively underscore that overexpressing the variable isoforms, as opposed to the C-type isoform C4, leads to a decrease in synaptic connectivity within the mouse neocortex.

### Combinatorial expression of γ-PCDHs regulates synaptic connectivity in the mouse neocortex

Our quest to understand the influence of combinatorial expression patterns of γ-PCDH isoforms on synapse formation led us to conduct a sequential *in utero* electroporation at E14.5 and E15.5 (Fig. 4A). In this intricate procedure, our objective was to deliberately manipulate the degree of similarity in expression between two distinct groups of neurons. To achieve this, we randomly selected isoforms, although with the notable exception of isoforms A5 and B8. These two isoforms were singled out due to their low expression levels as indicated by the single-cell sequencing data (Fig. 1D and Fig. 1-S1E). The various isoforms were thoughtfully combined to create similarity levels spanning from 0% (indicating no overlap, complete dissimilarity between the two groups) to 100% (representing complete overlap, indicating that the two groups are entirely identical) (Fig. 4B).

**Figure 4:**
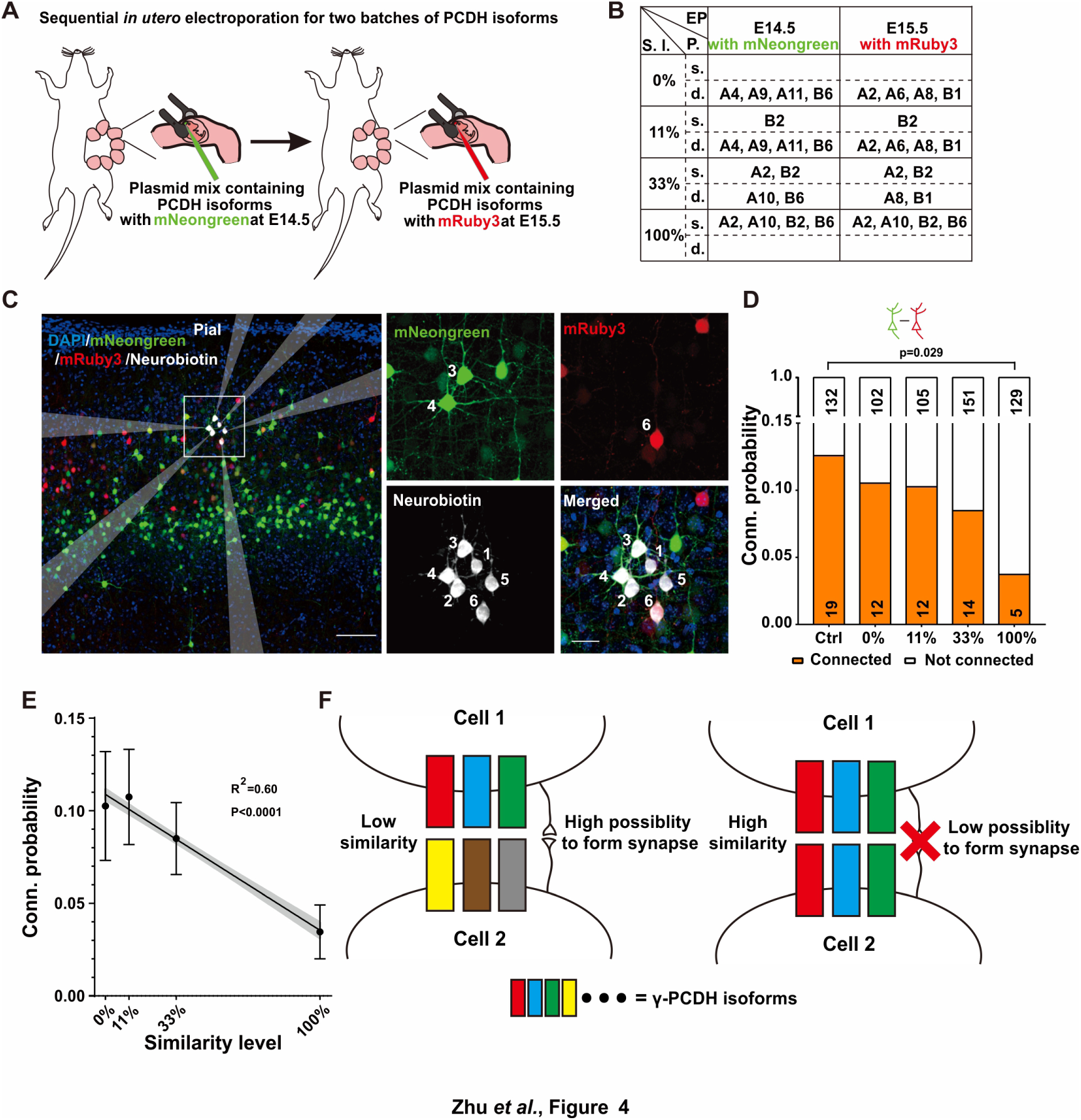
Diversified γ-PCDHs are critical for synapse formation in cortical neurons. (**A**) Diagram illustrating the sequential *in utero* electroporation process at E14.5 and E15.5. (**B**) Overview of the overexpressed γ-PCDH isoforms in different experiments, resulting in varying similarity levels between neurons. S.l.: Similarity level; EP: Electroporation; P.: plasmids mixture; s./d., Same or different isoforms in two electroporations. (**C**) Sample image of recorded neurons after two rounds of electroporation at E14.5 (mNeongreen) and E15.5 (mRuby3). Neurons labeled as positive for mNeongreen (cells 3 and 4), mRuby3 (cell 6), and negative without fluorescence (cells 1, 2, and 5). Neurobiotin was used in the internal solution to label recorded neurons. The translucent arrows show the positions of the electrodes. Scale bar, left: 100 μm, right: 25 μm. (**D**) Connectivity probability for neuron pairs overexpressing different sets of γ-PCDH isoforms (labeled with different fluorescence) following sequential in utero electroporation. Statistical differences were determined using the Chi-square test with FDR (Benjamini-Hochberg method) correction. (**E**) Correlation between the similarity level of overexpressed γ-PCDH combinations and the probability of synaptic connections. Each data point corresponds to the outcome of 100 bootstrapped samples derived from the source data presented in panel **D**. Error bars indicate the standard deviation (S.D) for each data point. The gray shaded area represents the 95% confidence interval of the curve fitting. (**F**) Graph summarizing the effect of γ-PCDHs on synapse formation.

For the purpose of electroporation at E14.5 and E15.5, we employed fluorescent proteins mNeongreen and mRuby3(Bajar et al., 2016) to tag isoforms, which allowed us to distinguish the cells electroporated on these respective days (Fig. 4A). Subsequently, whole-cell patch-clamp recordings were performed on layer 2/3 neurons in the neocortex using acute brain slices containing both green and red cells from P10-14 pups. Each set of recordings encompassed at least one mNeongreen^+^, one mRuby3^+^, and one nearby control neuron without fluorescence (Fig. 4C). The results were very clear: neurons sharing the same color displayed significantly lower connectivity (Fig. 4-S1A), aligning with our previous findings from single electroporation (as shown in Fig. 3E). In the group characterized by a 100% similarity level, the connectivity rate between neurons with different colors that were electroporated on different days was also significantly lower compared to the control (3.7%, 5/134 in the complete-overlap group; 12.5%, 19/151 in control pairs, 100% in Fig. 4D). However, as the similarity levels descended from 100% to 0%, the connectivity probabilities progressively reverted to the control level. The likelihood rebounded to 8.4% (14/165) for the pairs with a 33% similarity level and 10.3% (12/117) and 10.5% (12/114) for pairs with 11% or 0% similarity level, respectively (Fig. 4D). This compelling observation illuminated a fundamental principle: it is the similarity level of γ-PCDH isoforms shared between neurons, rather than the absolute expression of the protein within individual neurons, that dictates the regulation of synaptic formation.

Remarkably, in line with the single-electroporation experiment (gray bars in Fig. 3E), no significant changes were observed in the synaptic connectivity between electroporated neurons and nearby control neurons (Fig. 4-S1B). In summation, our findings illuminate a discernible negative correlation between the probability of synaptic connections and the similarity level of γ-PCDH isoforms expressed in neuron pairs (Fig. 4E). These discoveries underline the significance of the diversified combinatorial expression of γ-PCDH isoforms in regulating synapse formation between adjacent pyramidal cells. Simply put, the more similar the patterns of γ-PCDH isoforms expressed in neurons, the lower the probability of synapse formation between them (Fig. 4F).

## Discussion

Homophilic proteins cPCDHs are strong candidates for promoting synaptic specificity due to their combinatorial and stochastic expression pattern (Yagi, 2012, Kohmura et al., 1998, Toyoda et al., 2014). Our 5’-end single-cell sequencing data provided solid evidence for the combinatorial expression pattern of γ-PCDH isoforms in neocortical neurons. We further demonstrated the critical role of this diversity in synaptic connectivity through three lines of evidence. Firstly, the absence of γ-PCDH significantly increased functional connectivity between adjacent neocortical neurons. Secondly, electroporation-induced overexpression of identical γ-PCDH variable isoforms in developing neurons markedly decreased their connectivity. Lastly, using sequential *in utero* electroporation with different batches of isoforms, we found that increasing the similarity level of γ-PCDH variable isoforms expressed in neurons led to a reduction in their synaptic connectivity. These findings suggest that γ-PCDHs regulate the specificity of synapse formation by preventing synapse formation with specific cells, rather than by selectively choosing particular targets. It remains to be studied whether the diversified patterns of γ-PCDH isoforms expressed in different neurons have additional coding functions for neurons beyond their homophilic interaction.

Stochastic and combinatorial expression patterns of cPCDH have been identified in Purkinje cells(Esumi et al., 2005, Toyoda et al., 2014) and olfactory sensory neurons(Mountoufaris et al., 2017). However, it’s noteworthy that in serotonergic neurons, only one isoform, Pcdhac2, has been mainly detected (Chen et al., 2017). In our study, utilizing 5’-end single-cell sequencing, we have unveiled the stochastic and combinatorial expression patterns of variable γ-PCDH isoforms in neocortical neurons. These diverse observations across different cell types suggest that cPCDH diversity and the presence of ubiquitous C-type expression are not universal features throughout the brain(Kiefer et al., 2023). These distinct expression patterns of cPCDHs imply that this gene cluster might exert different roles in shaping neural connections in various brain regions. Furthermore, compared to the SMART seq results from the Allen Institute’s database and others(Lv et al., 2022) focusing on γ-PCDH, 5’-end single-cell sequencing used in our study not only detected more isoforms in individual cells but also revealed more neurons expressing C-type isoforms. The application of this approach may offer valuable insights for studying the functions of cPCDHs in a broader neurological context.

Previous studies by Molumby et al. demonstrated that neurons from the neocortex of *Pcdhg* knockout mice exhibited significantly more dendritic spines, while neurons overexpressing a single γ-PCDH isoforms had fewer dendritic spines in (Molumby et al., 2017). Our recordings are consistent with these previous morphology studies. Tarusawa et al. revealed that the absence of the whole cluster of cPCDH affected synaptic connections among lineage-related cells(Tarusawa et al., 2016). More recently, overexpression of the C-type γ-PCDH isoform C3 also showed a negative effect on synapse formation within a defined clone(Lv et al., 2022). In our study, we further demonstrated an increased synaptic connection rate between adjacent pyramidal neurons in the neocortex of *Pcdhg* knockout mice, while it decreased between neurons overexpressing single or multiple identical γ-PCDH variable isoforms. These effects were not just limited to lineage-dependent cells. Together with previous findings(Molumby et al., 2017, Tarusawa et al., 2016), our observations solidify the negative effect of γ-PCDHs on synapse formation among neocortical neurons. Some subtle differences exist between our findings and previous recordings(Tarusawa et al., 2016, Lv et al., 2022). Tarusawa et al. demonstrated that the connection probability between excitatory neurons lacking the entire cPCDH cluster in the layer 4 was approximately twofold higher at early stage P9-11, significantly lower at P13-16, and similar to control cells at P18-20 compared (Tarusawa et al., 2016). In our study, *Pcdhg*^-/-^ pyramidal neuronal pairs consistently exhibited a higher connection probability from P10 to P20. Two potential reasons could explain these differences. Firstly, we only removed γ-PCDHs instead of the entire cPCDH cluster, which includes α, β, and γ isoforms. Secondly, γ-PCDH might have different functions in neurons located in the layer 2/3 compared to the layer 4. Lv et al. found that overexpression of C-type γ-PCDH C3 decreased the preferential connection between sister cells(Lv et al., 2022). However, our study demonstrated that only variable, but not C-type isoform C4, had a negative impact on synapse formation. This discrepancy might be attributed to the lineage relationship, which could have an unknown impact on synapse formation. Building upon previous findings, it’s becoming increasingly evident that distinct C-type isoforms may play varied roles in shaping neural networks within the brain(Garrett et al., 2019, Steffen et al., 2023, Meltzer et al., 2023, Lv et al., 2022).

Since a single neuron can express multiple isoforms, deleting all γ-PCDH isoforms might mask the role of this combination. In this study, we manipulated the combinatorial expression patterns of γ-PCDH isoforms in nearby neocortical neurons through sequential *in utero* electroporation, expressing different batches of isoforms with adjustable similarities. We observed that when two neurons expressed identical variable isoforms (100% group), the likelihood of synapse formation between them was lowest. As the similarity level between two cells decreased, with fewer shared isoforms, the connectivity probability increased. The connectivity probability between neurons with different variable isoforms (0% group) did not differ from the control pairs (without overexpression). However, the connections between overexpressed and control neurons were not affected under both 100% and 0% similarity conditions. These observations suggest that the similarity level, rather than the absolute expression of the protein, affects synapse formation between neurons. While we observed a negative correlation between expression similarity and the probability of connectivity among neocortical neurons, further investigation is needed to determine the precise cellular mechanisms underpinning this correlation. It’s essential to explore whether this correlation arises directly from the formation of synapses or is a secondary effect resulting from cell positioning (Lv et al., 2022), synaptic pruning (Kostadinov and Sanes, 2015) or the influence of γ-PCDHs on the growth of axon/dendrite(Molumby et al., 2017, Molumby et al., 2016). Previous studies have established that the interplay between γ-PCDHs and neuroligin-1 plays a crucial role in the negative regulation of dendritic spine morphogenesis(Molumby et al., 2017). Consequently, exploring whether the similarity in the expression of γ-PCDHs between two neurons influences their interaction with neuroligin-1 could yield valuable insights.

Our findings also demonstrated that the overexpression of multiple γ-PCDH variable isoforms in one neuron only affected its connection if the other neuron overexpressed an identical combination of γ-PCDH isoforms. This highlights the pivotal role of the diversified combinatorial expression of γ-PCDHs of neurons in selecting their synaptic partners in the mouse neocortex. Notably, while the overexpression of the γ-PCDH C4 isoform had no discernible effect on synaptic connectivity, overexpressing six variable isoforms resulted in a reduced connection rate in *Pcdhg* cKO mice. These observations underscore the critical role of variable isoforms, as opposed to the C-type isoform C4, in synapse formation within the mouse neocortex. Although the overexpression of the γ-PCDH C3 isoform has been shown to have a negative effect on synapse formation between sister cells, but no effect on synapses among non-clone cells in the neocortex(Lv et al., 2022), the distinct functions of individual C-type isoforms require further thorough examination. It’s worth noting that our observations primarily stem from overexpression assays, providing insights into the effects of γ-PCDHs on synaptic connectivity. Exploring their impact under more physiological conditions using alternative approaches holds significant promise.

Furthermore, while the absence of γ-PCDHs causes significantly more synaptic formation among neocortical pyramidal neurons, evidence supports that their absence also leads to a significant reduction of synapse formation in other brain regions. For example, mice lacking γ-PCDHs exhibit fewer synapses in spinal cord interneurons(Weiner et al., 2005). Knocking down γ-PCDHs causes a decline in dendritic spines in cultured hippocampal neurons(Suo et al., 2012) and diminished astrocyte-neuron contacts in co-cultures from the developing spinal cord(Garrett and Weiner, 2009). The absence of γ-PCDHs leads to reduced dendritic arborization and dendritic spines in olfactory granule cells(Ledderose et al., 2013). Additionally, immuno-positive signals for γ-PCDHs are more frequently detected in mushroom spines than in thin spines(LaMassa et al., 2021). Moreover, our observations revealed that the absence of γ-PCDHs had a more pronounced impact on vertically aligned neurons than on horizontally situated pairs in the neocortex. These findings suggest that different mechanisms may be employed by synapses in different brain regions to achieve their specificity. Notably, different mechanisms have already been proposed for targeting specific inhibitory neural circuits in the neocortex, including “on-target” synapse formation for targeting apical dendrites and “off-target” synapse selective removal for somatic innervations(Gour et al., 2021).

In summary, our data demonstrate that the similarity level of γ-PCDH isoforms between neocortical neurons is critical for their synapse formation. Neurons expressing more similar γ-PCDH isoform patterns exhibit a lower probability of forming synapses with one another. This suggests that the presence of γ-PCDHs enables neocortical neurons to choose which neurons to avoid synapsing with, rather than selecting specific neurons to form synapses with. Whether there are specific attractive forces between cells to promote synaptic specificity remains an open question.

## Supporting information

Zhu-Supplementary data

## Acknowledgements

We thank Dr. Yifeng Zhang for providing the *Pcdhg flox/flox* mouse line and Dr. Zilong Qiu for *Nex-cre* mouse line, Dr. Jun Chu for sharing plasmids with mNeongreen and mRuby3, and other members in Xu lab for their discussions and technique supports. We are grateful to Prof. Mu-ming Poo and Song-hai Shi for critical reading of the manuscript. This work was supported by grants from the Training Program of the Major Research Plan of the National Natural Science Foundation of China, grant No. 91632101; Strategic Priority Research Program of the Chinese Academy of Sciences, grant No. XDB32010100; National Natural Science Foundation of China project 31671113; Shanghai Municipal Science and Technology Major Project, grant No. 2018SHZDZX05, the State Key Laboratory of Neuroscience and the Lingang Laboratory, grant No. LG-GG-202201-01.

## Author contributions

H. -T. X. conceived the project. Y. -J. Z. performed most recordings and single-cell analysis. C. -Y. D. and L. F. performed part of recordings. Y. -Q. W. prepared PCDH plasmids. H. Z. prepared mice. Y. -J. Z. and H. -T. X. wrote the manuscript.

## Notes

### Competing Interest Statement

The authors have declared no competing interest.

### Summary of Updates

More data was added for confirming the reduction of Alpha-PCDH in Pcdha CKO mice. More discussion was added.

