## Supplementary material for "Combinatorial expression of gamma-protocadherins regulates synaptic connectivity in the mouse neocortex": Zhu-Supplementary data

#### Materials and methods

##### Mice

All mice were handled in strict compliance with the approved protocol from the Animal Care and Use Committee of the Center for Excellence in Brain Science and Intelligence Technology/ Institute of Neuroscience, Chinese Academy of Sciences, and the committee at Lingang Laboratory. These mice were maintained in group housing conditions. *Pcdhg*$^{flox/flox}$, *Nex-cre* mice are maintained as heterozygotes within the C57BL/6 background. F2 mice resulting from the cross of these two lines contained *Pcdhg*$^{flox/flox}$/*Nex-cre* as experimental groups, and their wildtype littermates were used as controls for experiments. ICR mice from Vital River Laboratories were used in the overexpression experiments.

##### Single cell dissociation

At postnatal day 11 (P11), an ICR mouse was perfused with dissociation-artificial cerebrospinal fluid (aCSF) that had been thoroughly oxygenated with a 95% $O_2$/5% $CO_2$ mixture prior to use. The dissociation-aCSF used was based on choline-aCSF, consisting of the following components (in mM): 120 choline chloride, 2.6 KCl, 26 $NaHCO_3$, 1.25 $NaH_2PO_4$, 15 D-glucose, 1.3 ascorbate acid, 0.5 $CaCl_2$, and 7 $MgCl_2$ (with an osmolarity range of 300-310 mOsm), with the addition of 50 μM DL-APV (Abcam/ab120271) and 10 μM DNQX (Sigma/D0540).

The brain was promptly removed and immersed in dissociation-aCSF buffer at 0 °C. Half of the neocortex was stripped off and cut into small, approximately 100 μm diameter chunks. These chunks were then placed into a digestion buffer (dissociation-aCSF plus 100 units papain and 500 units DNase I) and allowed to incubate at 37°C for 60 min. Throughout the incubation, a gentle stream of 95% $O_2$/5% $CO_2$ was passed over the buffer. Following incubation, the sample was centrifuged at room temperature for 5 min at 300 g, and the resulting precipitate was collected and resuspended in 2 mL of Buffer II (dissociation-aCSF with 0.05% BSA).

The sample was then carefully triturated in sequence using fire-polished Pasteur pipettes with diameters of 600 μm, 400 μm, and 200 μm until no large, visible chunks remained. To remove debris, a debris removal solution (130-109-398, Miltenyi Biotec) was used in accordance with the manufacturer's protocol. Subsequently, the sample was resuspended with 500 μL Buffer II, which included 2.5 μg DAPI, and filtered through a 40 μm cell strainer before FACS sorting. Living cells were sorted via FACS based on their 350 nm fluorescence, ensuring a living cell rate of over 85%. The sorted cells were collected into Buffer II and subsequently subjected to centrifugation at 0°C and 500 g to obtain a single-cell suspension (refer to Fig. 1-S1).

**Library construction for 5' end single-cell sequencing**

We adapted the Mouse T Cell Chromium Single Cell V(D)J Enrichment Kit (PN-1000071, 10X Genomics) to create a library for the *Pcdhg* gene cluster. Although the kit was originally designed for sequencing T-cell receptor (TCR) genes in immune cells, we customized it by using specific primers for *Pcdhg* to sequence the 5' ends of different isoforms within this gene cluster.

We initiated the process by generating nanoliter-scale Gel Beads-in-emulsion (GEMs) using the Chromium Next GEM Chip G. Within these GEMs, we conducted reverse transcription, template switching, barcoding, and transcript extension (Fig. 1-S1B). The GEMs were later disrupted to facilitate cDNA amplification (Fig. 1-S1B). Upon completing the amplification, we divided the sample into two parts. One part was reserved for constructing an expression library for other genes at their 5' ends, while the other was used for constructing the enriched *Pcdhg* library (Fig. 1-S1C).

For the construction of the *Pcdhg* library, we replaced the original Reverse Outer & Inner Primers in Mouse T Cell Mix 1 and 2 with the *Pcdhg* nested-PCR primer mix, designed to target the constant regions of *Pcdhg*. This mix comprised the outer primer R1, situated within exon 4, and the inner primer R1, spanning exon 2 and 3 of the pcdhg gene. The complete sequences of the primers used can be found in the primer list at the end of this methods section.

Following the enrichment of *Pcdhg*, we adhered to the manufacturer's protocol for the remaining steps of library construction (left panel of Fig. 1-S1C). For the construction of the 5’ gene expression library, the amplified cDNA was immediately incorporated into the subsequent procedure in accordance with the kit's manual (right panel of Fig. 1-S1C). The two libraries were sequenced independently and subsequently intersected based on the cell barcodes (Fig. 1-S1D).

**Sequencing data profiling**

We obtained 212.66GB clean reads for the 5’ gene expression library and 183.30GB for the 5’ *pcdhg* expression library using the NovaSeq 6000 sequencing platform (Illumina, Novogene). After alignment with Cellranger v3.0.2 (10X Genomics) from the 5’ gene expression sequencing data, we identified 23753 cells (Fig. 1-S1D). These cells had an average of 29,843 reads per cell, and sequencing saturation reached 51.1%.

Cells were selected based on the criteria that feature RNA was within the range of 7,000 to 1,800, and mitochondrial RNA was less than 7%, resulting in 20419 cells (Fig. 1-S2A). Subsequently, 2981 potential doublets (15%) were removed using Doubletfinder, and the remaining cells were clustered using Seurat2 (resolution=2.4, optimized by the Clustree algorithm, Fig. 1-S1D and 1-S2B). After clustering, selected genes, including *Slc17a7, Neurod2, Neurod6, Slc32a1, Gad1, and Satb2* for neurons, and *Sox2, Car2, Ascl1, Aldh1l1, Pdgfra, Pdgfrb, Olig1, C1qa, and Dlx2* for non-neurons, were used to determine the identity of each cluster (Fig. 1-S2 to 4). Upon separating non-neurons from neurons, neurons were further divided into 15 clusters (cluster 0-14) (resolution=0.4, optimized by the Clustree algorithm, Fig. 1-S4A). Gene *Gad1, Gad2, Slc32a1, Slc17a1, Lamp5, Ndnf, Sncg, Vip, Sst, Pvalb, Cux2, Rorb, Fezf2, Sulf1, Foxp2, Mbp, Cldn5, Ctss,* and *C1qa* were used to identify the property of these clusters (Fig. 1-S4B). Cluster12 was classified as atypical neurons due to their high expression of *C1qa* and was excluded from further analysis.

To analyze the 5' *Pcdhg* expression sequencing data, we used Bowtie with the reference sequence of *Pcdhg* variable axons from all isoforms. Cell barcodes and UMI sequences were extracted from the aligned reads using a custom script in Matlab (R2018b, Mathworks). The *Pcdhg* expression matrix was generated from the intersection of data from the 5' gene expression library and the *Pcdhg* enrichment library (Fig. 1-S1D). In the output matrix, the *Pcdhg* expression level was represented by the number of UMIs. The *Pcdhg* expression matrix was filtered with a UMI count greater than 1 and binarized for subsequent analysis (Fig. 1-S1D). Various UMI thresholds for the total *Pcdhg* were applied to assess cell fractions for all isoforms in individual cells (Fig. 1D and Fig. 1-S1E). A UMI count of greater than 10 was chosen for the similarity level analysis in 2834 neurons to ensure the quality of the assay.

**Definition of the Similarity level**

The similarity level was computed using the formula $\frac{A \cap B}{A \cup B}$, where A and B represent the numbers of γ-PCDH variable isoforms (excluding C-type isoforms C3, C4, and C5) expressed in two compared cells; $A \cap B$ represents the number of isoforms shared in both cells and $A \cup B$ equals to the total number of isoforms expressed in these two cells.

In fig. 1-S5, we also used the Euclidean distance and the Normalized Euclidean distance to measure the similarity between two neurons. The Euclidean distance is a widely used method for describing the spatial distance between two nodes in a multi-dimensional space. In our dataset, each neuron represents one node in a 19-dimensional space, with each axis representing one variable isoform. The coordinate of each node in each axis is determined by the expression level of the corresponding isoform (counted by UMI). If a cell pair has a lower Euclidean distance, then these two cells express a more similar combination of γ-PCDH isoforms.

The Euclidean distance is calculated as follows:

$$\text{Euclidean distance} = \sqrt{\sum \left( {\Delta_{UMIcounts(pcdhga1)}}^2 + {\Delta_{UMIcounts(pcdhga2)}}^2 + \cdots + {\Delta_{UMIcounts(pcdhgb8)}}^2 \right)}$$

However, if two cells have a substantial difference in total expression levels, the Euclidean distance may not be sufficiently sensitive. Therefore, we also explored the "Normalized Euclidean distance" to better assess the dissimilarity between two neurons. After normalizing the total expression level of variable isoforms (19 in total) in each cell to 1, we calculated the Normalized Euclidean distance. The Normalized Euclidean distance falls within the range of [0, √2].

**Co-occurrence**

We adopted a concept from Wada's study to investigate the potential interactions among different isoform expressions at the population level. Co-occurrence analysis was used to determine whether each isoform expresses independently.

To perform this analysis, we first generated an expected matrix by shuffling data 100 times, drawing from different clusters (or the total population), assuming that all the isoforms were expressed randomly based on their expression frequencies. Subsequently, we plotted the fraction of cells with various numbers of isoform types from both the observed expression matrix (observed data) and the expected matrix (shuffled data) for the entire population (Fig. 1F and Fig. 1-S7).

We then computed the variance of this distribution for cells in different clusters or the total population. The expected variance was determined using the following formula, taking into account the expression probability of each isoform (pi):

$$\sigma_{expected}^2 = \sum_{i=1}^{N_{isoforms}} p_i(1 - p_i)$$

Subsequently, we used a Z-test to evaluate the statistical difference between the observed variance and the expected variance. If the observed variance was found to be higher than the expected variance, it indicated that the genes were co-occurring.

**Generation of *pcdha* cKO mice**

*Pcdha* conditional knockout (cKO) mice were generated using the CRISPR/Cas9 approach by Biocytogen Pharmaceuticals (Beijing). Candidate sgRNAs for intron 1 and intron 2 of *Pcdha1* were designed using the CRISPR design tool (http://crispr.mit.edu) and screened for on-target activity using the Universal CRISPR Activity Assay kit (UCATM, Biocytogen Pharmaceuticals (Beijing) Co., Ltd).

To minimize random integrations, a circular donor vector was employed. The gene-targeting template contained a 5' homologous arm, the target fragment (exon2), and a 3' homologous arm. This templated was used to repair the double-strand breaks generated by Cas9/sgRNA. Cas9 mRNA, the template, and sgRNAs were co-injected into the cytoplasm of one-cell stage fertilized C57BL/6 egg. These injected eggs were then implanted into the oviducts of Kunming pseudopregnant females to produce F0 mice.

F0 mice with the correct genotype were bred with C57BL/6 mice to establish germline-transmitted F1 heterozygous mice. The PCR-positive F1 mice were further confirmed with Southern blot analysis (Fig. 2-S2). DNA sequencing was used to confirm the absence of off-target effects (Fig. 2-S3-4). Exon 2 of *Pcdha genes* will be removed with the assistance of Cre, resulting in an 813 aa (796aa native, 17aa in-frame shifted) truncated protein that may be subject to nonsense-mediated decay.

**Southern blot**

The digoxin (DIG) labeled probes were synthesized through PCR, using the PCR DIG Probe Synthesis kit (11636090910, Roche), with genomic DNA extracted from mouse tails and digested with BamHI or StuI (R0136, R0187, New England BioLabs). The DNA was separated on a 1% agarose gel and transferred to a positively charged nylon membrane (HyBond N+, Amersham). The hybridization process was carried out at 42℃ overnight using the DIG Easy Hyb Granules (11796895001, Roche). Following hybridization, the signals were subsequently amplified using the DIG Luminescent Detection kit (11363514910, Roche).

**SgRNA off-target test**

Potential sgRNA off-target sites were identified using the website https://crispr.cos.uni-heidelberg.de/. Subsequently, primers (available in the provided list) were designed based on the website https://crispr.bme.gatech.edu/ to target the top 10 off-target regions, aiming for a product size of approximately 600 bp.

For the experimental process, male *pcdha*$^{+/flox}$ mice were bred with female *pcdha*$^{+/flox}$ mice. From the resulting offspring, F1 generation homozygotes, heterozygote littermates, and WT littermates were each selected for genome DNA extraction using the Mouse direct PCR Kit (B40018, Bimake). Following DNA extraction, PCR amplification was carried out, and the resulting PCR products were subsequently subjected to sequencing (Sangon, Shanghai).

**Acute slice preparation**

The brains of mice aged between P9 and P14 were extracted immediately after sacrifice and placed on ice. Coronal sections with a thickness of 300 μm were prepared using a vibratome (VT1200S, Leica). The sections were cut in choline-modified artificial cerebrospinal fluid (choline-aCSF) containing the following concentrations (in mM) 120 choline chloride, 2.6 KCl, 26 $NaHCO_3$,1.25 $NaH_2PO_4$, 15 D-glucose, 1.3 ascorbate acid, 0.5 $CaCl_2$, and 7 $MgCl_2$. This solution maintained an osmolarity of 300-310 mOsm. The slices were then transferred to a chamber with normal aCSF containing (in mM) 126 NaCl, 3 KCl, 26 $NaHCO_3$, 1.2 $NaH_2PO_4$, 10 D-Glucose, 2.4 $CaCl_2$, and 1.3 $MgCl_2$ (with an osmolarity of 300-310mOsm) and incubated at 32°C for 30 min. Afterwards, they were maintained at room temperature until recording.

For mice older than P14, pentobarbital sodium was administered for anesthesia via intraperitoneal injection (50-90 mg/kg). Following anesthesia, the brains were then perfused with 0°C NMDG-aCSF containing (in mM) 93 N-Methyl-D-glucamine, 2.5 KCl, 1.25 $NaH_2PO_4$, 30 $NaHCO_3$, 25 D-glucose, 20 HEPES, 5 Na-ascorbate, 3 Na-pyruvate, 2 Thiourea, 10 $MgSO_4$, 0.5 $CaCl_2$ and 12 N-Acetyl-L-cysteine (with a pH of 7.3-7.4 and an osmolarity of 300-310 mOsm). Subsequently, the brains were coronally sectioned, following the same procedure as described above. The slices were incubated in NMDG-aCSF at 37°C for 12-15 minutes, then transferred to normal aCSF, and kept at room temperature for one hour before recording.

It's worth noting that all of the solutions mentioned were continuously oxygenated with a mixture of 95% $O_2$/5% $CO_2$.

**Electrophysiology**

During recording, the slices were consistently perfused with normal aCSF maintained at 30°C and oxygenated with a mix of 95% $O_2$ and 5% $CO_2$. Cell visualization within the slices was performed using a differential interference contrast microscope

(BX51W1, Olympus) equipped with both a 5X/NA 0.1 objective and a 40X/NA 0.8 W objective.

Glass pipettes, with resistance at 8-12 MΩ, were prepared with a Flaming/Brown micropipette puller (P-97, Sutter Instrument). The internal pipettes solution contains (in mM) 126 potassium gluconate, 2 KCl, 2 $MgCl_2$, 10 HEPES, 0.2 EGTA, 4 $Na_2$-ATP, 0.4 $Na_3$-GTP, 10 creatine phosphate and 10 ng/ml Alexa Fluor 405 Cadaverine. Multiple whole-cell patch-clamp recordings, involving up to 6 channels, were conducted using amplifiers (Multiclamp 700B, Molecular Devices) and Digidata (Digidata 1550A, Molecular Devices). Subsequently, the data were analyzed using pClamp9 (Molecular Devices).

Electrophysiological data underwent low-pass Bessel filtering at 2 kHz and were digitized at 20 kHz. The feedback resistor was set at 5 GΩ. All the recordings had a series resistance less than 50 MΩ. Neuronal connectivity was assessed by injecting current to one of the recorded cells under the current-clamp mode to evoke action potentials. Simultaneously, evoked excitatory postsynaptic currents (EPSC) in other receiver cells were examined under the voltage-clamp mode, maintaining a holding potential of -70 mV.

To determine the presence of meaningful EPSC, the size of currents in receiver cells was quantified one second before stimulation to obtain the average and standard deviation (SD). A current within 3 mini-second after stimulation that was 3 times of SD greater than the average and exhibited minimal jittering among different trials (shorter than 0.2 ms), was considered a meaningful EPSC. Each neuron pair in every recording was tested at least ten times in both directions.

##### ***In utero* electroporation**

For pregnant ICR mice or pcdhg cKO mice, the date of plug detection was designated as embryonic day 0.5 (E0.5). At the gestational stages of E14.5 or E15.5, the pregnant mice were anesthetized using 5% isoflurane and subsequently maintained under 1.0% isoflurane with a gas flow rate of 0.8-1 L/min. Following a surgical incision, the uterine horns were exposed, and a plasmid mixture with a total concentration of 5 mg/ml plasmid (equal molar mixture if there are multiple plasmids) was injected into the embryonic lateral ventricle using a glass pipette. This plasmid mixture included 1 mg/ml Fast green (Sigma). After the injection, the utero was clamped with electrodes with a diameter of 7 mm (45-0118, Tweezertrodes), and five 30 V pulses of 50 ms duration were delivered with 1s intervals.

In sequential *in utero* electroporation experiment, surgeries were performed at E14.5 and E15.5, and the plasmids were injected into the same lateral ventricle and delivered at a similar angle. Following the procedure, lincomycin hydrochloride and lidocaine hydrochloride gel were applied to the wound after stitching it up.

**qRT-PCR**

For the confirmation of *pcdhg* overexpression (Fig.3A, B): The brains of mice electroporated at P11-14 were promptly removed at sacrifice and placed into 4°C choline-aCSF. The specific target region was then identified using a fluorescent microscope, and approximately 500 μm-thick coronal slices were prepared. Tissue blocks containing the fluorescently labeled region, as well as the contralateral control side, were extracted from the same coronal slice and immediately transferred into an RNA extraction buffer.

For *Pcdha* knockout confirmation (Fig.2-S2D), the semi-cortex of P11 mice was isolated and transferred into an RNA extraction buffer. Tissue RNA was extracted using the Total RNA extraction Kit (R4011-02, Magen) and then reverse transcribed using the PrimeScript™ RT Master Mix (RR036A, Takara).

Primers, the sequences of which can be found in the primers list, were designed with the assistance of PrimerBlast (NCBI). SYBR Green Real-time PCR Master Mix (QPK-201, Toyobo) and the LightCycler 480 II (Roche) were employed for qPCR.

The expression level was calculated as $2^{Ct(Pcdhg)-Ct(GAPDH)}$, and the normalized expression level (Fig. 3B) was determined by dividing the expression level of the target fluorescent region by the expression level of the contralateral control region.

**Estimating the numbers of overexpressed *pcdhg* isoform**

Five plasmids, each containing one of the following isoforms *pcdhga*2, *pcdhga8*, *pcdhga10*, *pcdhgb1*, and *pcdhgb2*, tagged with mNeongreen were equimolarly mixed. A sixth plasmid, pCAGGS-*pcdhga6*-p2A-mCherry, was included for electroporation at E15.5. The mice were sacrificed at P7, and their brains were sectioned coronally with a vibratome at a thickness of 60 µm. The mCherry signal was enhanced by using an anti-RFP antibody (Rockland 600-401-379) to achieve a brightness similar to mNeongreen. The images were captured using a confocal microscope (Nikon C2, 20X, NA 0.75, Fig. 3-S1a).

The co-transfection rate was calculated by the following formula. For simplicity, we'll use two plasmids first as an example.

When two plasmids, one containing GFP and the other RFP, were used for an electroporation at a 1:1 molar ratio:

$$R_{GFP-only} = \left(\frac{1}{2}\right)^n \quad (1)$$

$$R_{RFP-only} = \left(\frac{1}{2}\right)^n \quad (2)$$

$$R_{co} = 1 - \left(\frac{1}{2}\right)^n - \left(\frac{1}{2}\right)^n = 1 - \left(\frac{1}{2}\right)^{n-1} \quad (3)$$

The percentages of cells that only express one fluorescent protein in all transfected cells (the percentage of green or red cells in all transfected cells) are represented by $R_{GFP\text{-}only}$ and $R_{RFP\text{-}only}$, respectively. The percentages of cells that co-express both fluorescent proteins (the percentage of yellow cells in all transfected cells) is represented by $R_{co}$. The variable 'n' in the formula is the number of received plasmid for each cell.

When 6 plasmids, 5 tagged with GFP and one with RFP, were electroporated with equal molar ratio:

$$R_{GFP-only} = \left(\frac{5}{6}\right)^n \quad (4)$$

$$R_{RFP-only} = \left(\frac{1}{6}\right)^n \quad (5)$$

$$R_{co} = 1 - \left(\frac{5}{6}\right)^n - \left(\frac{1}{6}\right)^n \quad (6)$$

$$R_{Red\ in\ total} = R_{co} + R_{RFP-only} = 1 - \left(\frac{5}{6}\right)^n \quad (7)$$

Note that formula (1)-(7) only describe the most idealized situation where 'n' is a fixed number for every cell. However, when these formulas were applied to fit our data, they did not provide a good fit. We recognized that 'n' was not a fixed number for every cell, and we hypothesized that 'n' might obey a normal distribution ($n \sim N(\mu, \sigma^2)$). A normal distribution is characterized by two parameters: the mean (μ) and the standard error (σ). Therefore, we modified functions (5) and (7) to incorporate the idea that 'n' follows a normal distribution.

$$R_{RFP-only} = \frac{\sum_{n=1}^{k} \left(\frac{1}{6}\right)^n}{k}, \quad n \sim N(\mu, \sigma^2) \qquad (8)$$

$$R_{Red\ in\ total} = \frac{\sum_{n=1}^{k} 1-\left(\frac{5}{6}\right)^n}{k}, \quad n \sim N(\mu, \sigma^2) \qquad (9)$$

The parameter 'k' in the formulas represents the total number of affected cells. We simulated 'n' by systematically varying the mean (μ) and standard error (σ) with integer values and found that the mean value of 18 (μ=18) and the standard error of 6 (σ=6) closely matched the experimental data (Fig. 3-S1C). We then simulated 10,000 cells following this distribution and counted the number of expressed isoform types in each cell. Ultimately, we found that each affected neuron expressed an average of 5.6 isoform types (Fig. 3-S1D).

**Single-cell RT-PCR**

For single-cell RT-PCR, we used the SuperScript II Kit (18064014, Invitrogen). Neurons were collected from acute brain slices using a glass pipette with a resistance of 3-4 MΩ, pre-filled with approximately 1 μL of aCSF. The cell contents were then gently expelled into a PCR tube containing 0.5 μL Rnaseout (10777019, Invitrogen), and the samples were promptly frozen with liquid nitrogen.

Subsequently, RT mix1 (~1 μL sample, 0.5 μL Rnaseout, 10 mM dNTP, 1 μL primer mix (GR1 and β-actin R1, 2.5 μM), 1.5 μL $ddH_2O$) was added to the sample tubes and kept at 65°C for 5 min. The tubes were then quickly chilled on ice. RT mix2 (1 μL 10X RT buffer, 2 μL 25 mM $MgCl_2$, 1 μL 0.1 M DTT, 0.5 μL Rnaseout, and 0.5 μL SuperScript II) was added to the sample/mix1 for the reverse transcription, which took place at 50°C for 60 min, followed by 85°C for 5 min.

For the nested PCR, we employed LATaq Kit (RR002B, TaKaRa). The primer mix for the first round of the PCR contained forward primers *Gb* F1a, *Gb* F1b, *Ga* F1a, *Ga* F1b, *Ga* F1c, *Ga* F1d， and reverse primer *GR*1. Another primer mix, consisting of a forward primer corresponding to each isoform and reverse primer *GR*2, was used for the second round of PCR. The sequences of each primer are listed in the following primers list. Due to the high sensitivity achieved after two rounds of PCR, faint bands appeared in some negative controls, leading to potential false-positive signals (indicated by red stars in Fig. 3D). A true positive signal was defined as a band with at least five times higher intensity than the false-positive band.

#### Bootstrap resampling

We employed bootstrap resampling using the 'bootstrp' function in Matlab (R2018b, Mathworks), setting the 'nboot' parameter to 100.

#### Primers list

| Primer name | Sequence (5'-3') | Working conc.(μM) |
|---|---|---|
| | **Library construction for 5' end single-cell sequencing** | |
| Outer R1 | GTAAACTGGGGTCCGTATC | 1 |
| Inner R1 | CATTTTGGGATCCGCTCGT | 0.5 |
| | **Generation of *pcdha* cKO mice** | |
| Southern probe A | CGGGGGTAAGTATAGTTTGAACTTTGAACCCATT<br>GCCTGGCACTGCAACCGCTTTATGTTGCTATGATC<br>ATTGTAACCTAGAAGAAATCCCCCCTTCCATTTCC<br>AGTTTAGTATCCATGCCTGATTTAGTCTCTAGTGA<br>TTCCGAGTACCTTCTGGCTCCCCACCAGTCTTCAG<br>GTGCCATATTTTCTTCTTTATCAATTCTACTTTGGA<br>ATGACAGTTTGTGGGCAGAGGGCACACAAATTTT<br>CTTTCTAGGACACAGGTGGTGCAGAAAGAGTTGG<br>AGTCCTTGTGGATTTCATCTACTTCCTGCTAAGCA<br>ATCAGAGCCTAGTGAATCAGGAGGTAAAGCATGA<br>GAAAGGGGCAGGGAGCCAGTGCAGATTTACAAGT<br>AGGTTTGGGTAGCCAAACA<br>CTGACTTGAATCAGACTTCCTTTGCATATTATAAT<br>ATAGCCTTAATTAATTTGATTAGCATAATAATATT<br>TTGATCATTATTGAACAAGTAGCAGTGGTTTAAA<br>AATTCAGAGGTCAAGAGAACTCTTTAGCTATGTTT | |

| | | |
|---|---|---|
| | CAAAAACAGGTGTTCTTAAAAGTGGCTTTTACAG | |
| Southern probe B | GACCCCACTTGGATCATAGTTTCTATGAATATTAA | |
| | ATATGAACATAATTCACCTGATTAATATAATAAC | |
| | ATCTTGACCAATATTTAAACCAGTAGCCATGATTT | |
| | TAAAATTTATAAGTCAAGAGAATTCTTTAACAAT | |
| | GCTTTAAATTTGTTAACCTAAAAATTATACAAGCT | |
| | CTTTCAAGAATTTGAACTTGGCACATGCAAAGAA | |
| | ATATCTGACATTGCAAAGGTGGAGAGA | |
| Target1_1F | AGTTTGCAGACACCTGTTGGAACC | 0.25 |
| Target1_1R | CTCCTTACTTACCACAGGGACTGA | 0.25 |
| Target1_2F | GTAATGGTGGTGCAGGCCCTA | 0.25 |
| Target1_2R | GCAGCCTCAGAGGAAGTGCA | 0.25 |
| Target1_3F | CCAAATGGGGGAAAGCCCAG | 0.25 |
| Target1_3R | GGAGGTCAGAGGTCAATTCATGG | 0.25 |
| Target1_4F | AGCAAGCAAGAAGGGTCAAGGACA | 0.25 |
| Target1_4R | AACCCAGACATTGCCCTTGGTTG | 0.25 |
| Target1_5F | GCTGGATGTTTTGACTCTGTTGGG | 0.25 |
| Target1_5R | GAGCCAGAAACTGGGAACAACC | 0.25 |
| Target1_6F | GCTTTCACATCCATGTCACAGTCC | 0.25 |
| Target1_6R | GCTGGCATTTCTCCTTATGCTTCC | 0.25 |
| Target1_7F | GAACATTCTCCCCAAGTGTCAAGG | 0.25 |
| Target1_7R | AGAACCTCTAGAAAGGTCCCAGTC | 0.25 |
| Target1_8F | GCTGGGGTAGTCACATCTAGATC | 0.25 |
| Target1_8R | TTACCAATGGCCTGGCCTTGGCT | 0.25 |
| Target1_9F | TCAAGGCAGAGGAGAAGGGC | 0.25 |
| Target1_9R | GGCTTTCCAGTCTTCACAACCATC | 0.25 |
| Target1_10F | GTGGTCAATGTCAGTCAATGTCCC | 0.25 |
| Target1_10R | AGGATACTTGAGGGTACCCAAGG | 0.25 |
| Target2_1F | GGCAGAGGTCATCACTGAGTTC | 0.25 |
| Target2_1R | CCTAGCTACTTCCTAGCAACAGTC | 0.25 |
| Target2_2F | GGGTTTGTGAAACCTCAGATCTGC | 0.25 |
| Target2_2R | CCTAGGCTATCCAAGGCCTC | 0.25 |
| Target2_3F | ACCTGGCACACAGATTTGGATGAC | 0.25 |
| Target2_3R | GCCTAATGGCCTGGGTTTTATCC | 0.25 |
| Target2_4F | GTGGCTGCCAGTCTGACTGT | 0.25 |
| Target2_4R | CGCTTCTCCCATTGGTGAAGTG | 0.25 |
| Target2_5F | GACTGTAATGTCTGCGGTAACTGC | 0.25 |
| Target2_5R | GACCAAGTCAGGGGCAGCTTA | 0.25 |
| Target2_6F | GCTCCTTTTGGATTGTCCCTTCCT | 0.25 |
| Target2_6R | GGACCCTCTTGATGCCTCTATG | 0.25 |
| Target2_7F | AGACCTCAGCTCCCTTAATACTGC | 0.25 |
| Target2_7R | GGTTTTGCCCCAGCAGGTGA | 0.25 |
| Target2_8F | AACGGGTTCACTGCTCTGTGG | 0.25 |
| Target2_8R | GTCACCCTAGTTAGGACAAAGCAG | 0.25 |

| | | |
|---|---|---|
| Target2_9F | GATTTTGAGTGTCAGGCAAGCGG | 0.25 |
| Target2_9R | GCAGGAAGGATCAGAAGTTCAAGG | 0.25 |
| Target2_10F | TCCATCTCTTCTTCTCGGTAAGGC | 0.25 |
| Target2_10R | CTCAGAACTCAGGAGGCAGG | 0.25 |
| | **qRT-PCR** | |
| *Ga2* qPCR F | GATGCGGACGTTGGAGAGAA | 0.5 |
| *Ga2* qPCR R | CGTCAGAGGCGACTAGAACC | 0.5 |
| *Ga8* qPCR F | TCAGCGTAGGACAGATTCGC | 0.5 |
| *Ga8* qPCR R | CGACCTACCTCTGGAGACGA | 0.5 |
| *Ga9* qPCR F | TCAGTTGAGCCCAAGTTTCCT | 0.5 |
| *Ga9* qPCR R | TCACCATTTTGGGATCCGCT | 0.5 |
| *Ga10* qPCR F | AATCTCTCAGAGCGCACCAAA | 0.5 |
| *Ga10* qPCR R | CCTACCTCTGGAGATGATGCG | 0.5 |
| *Ga11* qPCR F | AGACTCAACTACAGTTTTAGGCA | 0.5 |
| *Ga11* qPCR R | TGGGTGTGAGCGTTATTCCG | 0.5 |
| *Gb1* qPCR F | ATGAGGGGACATTACCCTATTCC | 0.5 |
| *Gb1* qPCR R | GCGGGGCTTGCTGGAAA | 0.5 |
| *Gb2* qPCR F | TCAACCTGGCCTCAGCTCTA | 0.5 |
| *Gb2* qPCR R | CTGGTACCCAAGAGTCGTCG | 0.5 |
| *Gb6* qPCR F | TCGTTTCCGGTAGTTCTCCTG | 0.5 |
| *Gb6* qPCR R | GGCTTGAGAGAAACGCCAGT | 0.5 |
| *GC3* qPCR F | AAGCGCTAACCCGCTGAAAG | 0.5 |
| *GC3* qPCR R | GCAGAAGCAAAACTCCCACC | 0.5 |
| *GC4* qPCR F | TCCACGATGATAACGAGCCC | 0.5 |
| *GC4* qPCR R | CGTCGTAGCATTGGGAGGAA | 0.5 |
| *Pcdha* qPCR F | TACTCTGCCTCGCTAAGAGC | 0.5 |
| *Pcdha* qPCR R | TGGTGTTGCACTGGATACTG | 0.5 |
| | **Single-cell RT-PCR** | |
| *Gb* F1a | GACACAATGCCTGGCTGTC | 0.13 |
| *Gb* F1b | GACACAATGCCTGGCTATC | 0.05 |
| *GR1* (also as RT primer) | TCGTTTGCCAGCGGCATTG | 0.5 |
| *Gb1* f2 | CACATTCTACGACTATGGCAGG | 0.1 |
| *Gb2* f2 | ACCAGGTACTCTTGGAGACAC | 0.1 |
| *Gb4* f2 | CTACAATTTGCAGATTCAGCG | 0.1 |
| *Gb5* f2 | ATGGTGCACATGCTGGAAAGAC | 0.1 |
| *Gb6* f2 | CGTTGGGCAAACGGGAAAGG | 0.1 |
| *Gb7* f2 | GCCGGGAGATCAACTTAAAC | 0.1 |
| *Gb8* f2 | GAGTGTCGCTGAGGAGAATA | 0.1 |
| *GR2* | ATTGGTCAGCGTGGCATTGCT | 0.1 |
| *Ga* F1a | CCTCTCTCCGCCACTGTCA | 0.10 |
| *Ga* F1b | TCTACCTGGTGGTGGCAGT | 0.05 |
| *Ga* F1c | TGGTGGCAGTGGACAGAGA | 0.05 |

| | | |
|---|---|---|
| *Ga* F1d | TGGTGGCAGTGGACAAAGA | 0.10 |
| *Ga1* f2 | CAGGAATTTTTGTCAGCACC | 0.1 |
| *Ga2* f2 | CTGATTTCCTCTCAGCACCTC | 0.1 |
| *Ga3* f2 | ACGAAAGAAGACCCCACGCTG | 0.1 |
| *Ga4* f2 | ATAAAAGGAGACTCCAGTCTGCAG | 0.1 |
| *Ga5* f2 | CACAAAGAAGAGCCCGGAGATG | 0.1 |
| *Ga6* f2 | GAAAGGTGTGTCAAATATGTCC | 0.1 |
| *Ga7* f2 | TTCAGGTGGTGGCCTGGAAGAT | 0.1 |
| *Ga8* f2 | GATGAAGATGCTTGCGCTCCG | 0.1 |
| *Ga9* f2 | GCAGTTCAGTTGAGCCCAAGT | 0.1 |
| *Ga10* f2 | CAAGTGTCCTGTAGAAGACG | 0.1 |
| *Ga11* f2 | GACTCAACTACAGTTTTAGGC | 0.1 |
| *Ga12* f2 | GGCAGATCTAGACAATCTAG | 0.1 |
| *β-actin* F | GTCGTACCACAGGCATTGTGATGG | 0.5 |
| *β-actin* R1 | CTAGAAGCACTTGCGGTGC | 0.5 |
| *β-actin* R2 | GCAATGCCTGGGTACATGGTGG | 0.1 |

### Supplementary figures:

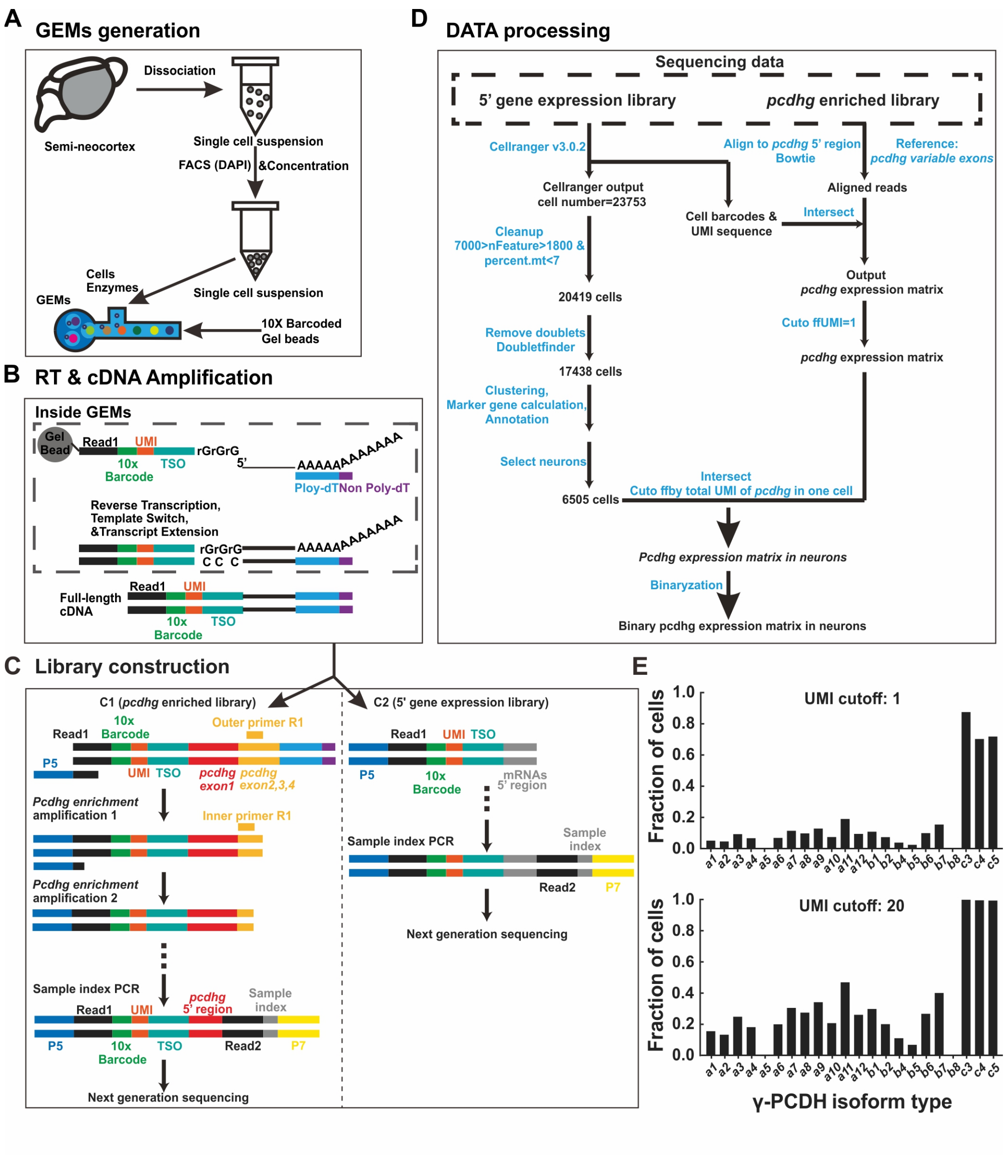


Zhu *et al.*, Figure 1-Figure supplement 1

**Figure 1-S1: Overview of the 5'-end single-cell sequencing procedure for *pcdhg* isoforms.** (**A-D**) Schematic representation of the 5'-end single-cell sequencing protocol. For more detailed information, please refer to the methods section. Key components in (**C**) and (**D**) are outlined in the legend below. P5/P7: Illumina sequencing priming sites; Read1: Illumina R1 sequence; 10x Barcode: a 16-nucleotides (nt) cell barcode; UMI: a 10-nt unique molecular identifier; TSO: a 13-nt template switch oligo; Read2: an Illumina R2 sequence; Sample index: an 8-nt sequence for sample identification. (**E**) The proportion of neocortical neurons expressing different *Pcdhg* isoforms, analyzed under varying UMI cutoff 1 (top) or 20 (bottom).

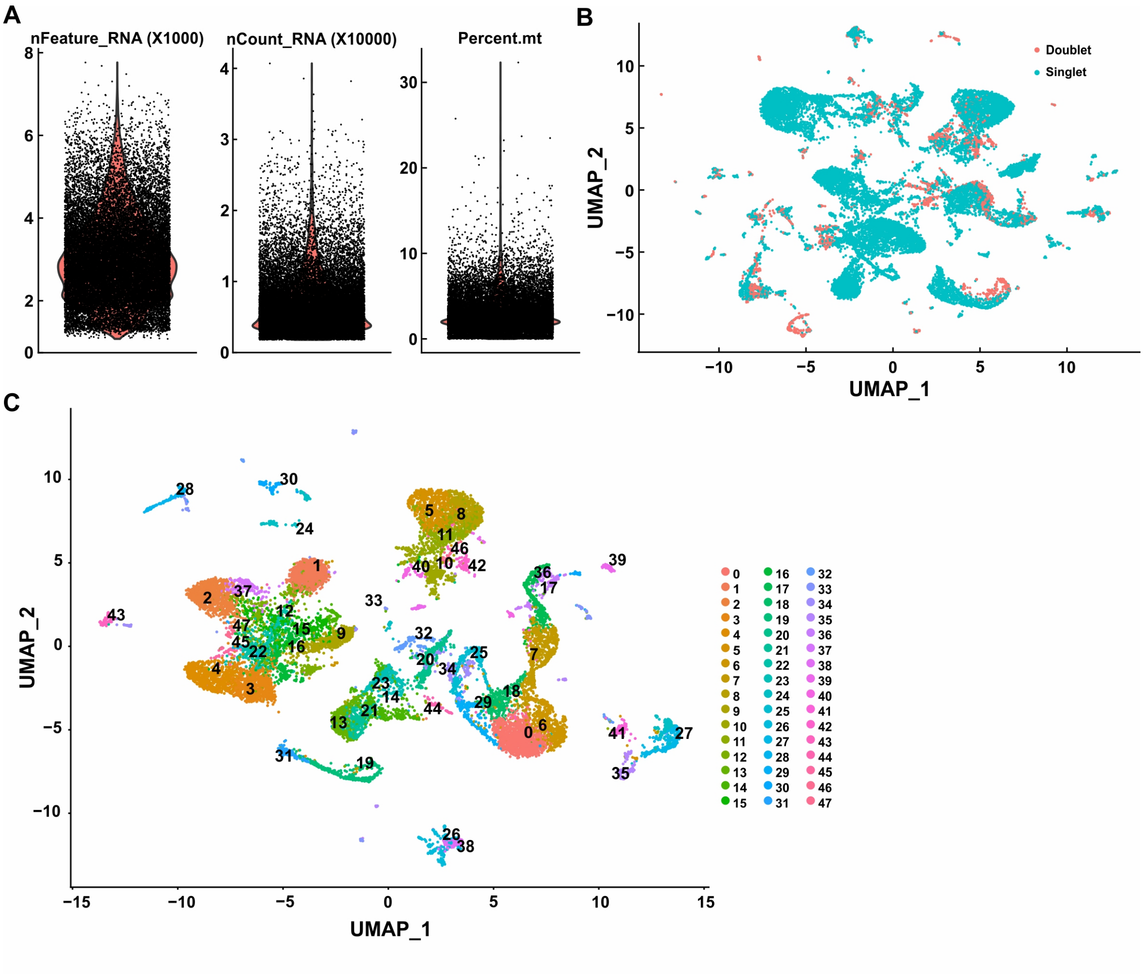


Zhu *et al.*, Figure 1-supplement 2

**Figure 1-S2: Single-cell RNA sequencing data profiling.**

(**A**) Quality control for sequenced cells. Cells were selected for analysis by the range of 7000> nFeature_RNA>1800, and mitochondrial RNA< 7%. nFeature_RNA: total feature (gene) counts per cell; nCount_RNA: total counts number per cell; percent.mt: the percentage of mitochondrial RNA per cell. (**B**) UMAP analysis of the entire cell dataset (n=20419), red dots: potential doublets (n=2981); green dots: singlet (n=17438). (**C**) UMAP analysis of singlets (n=17438).

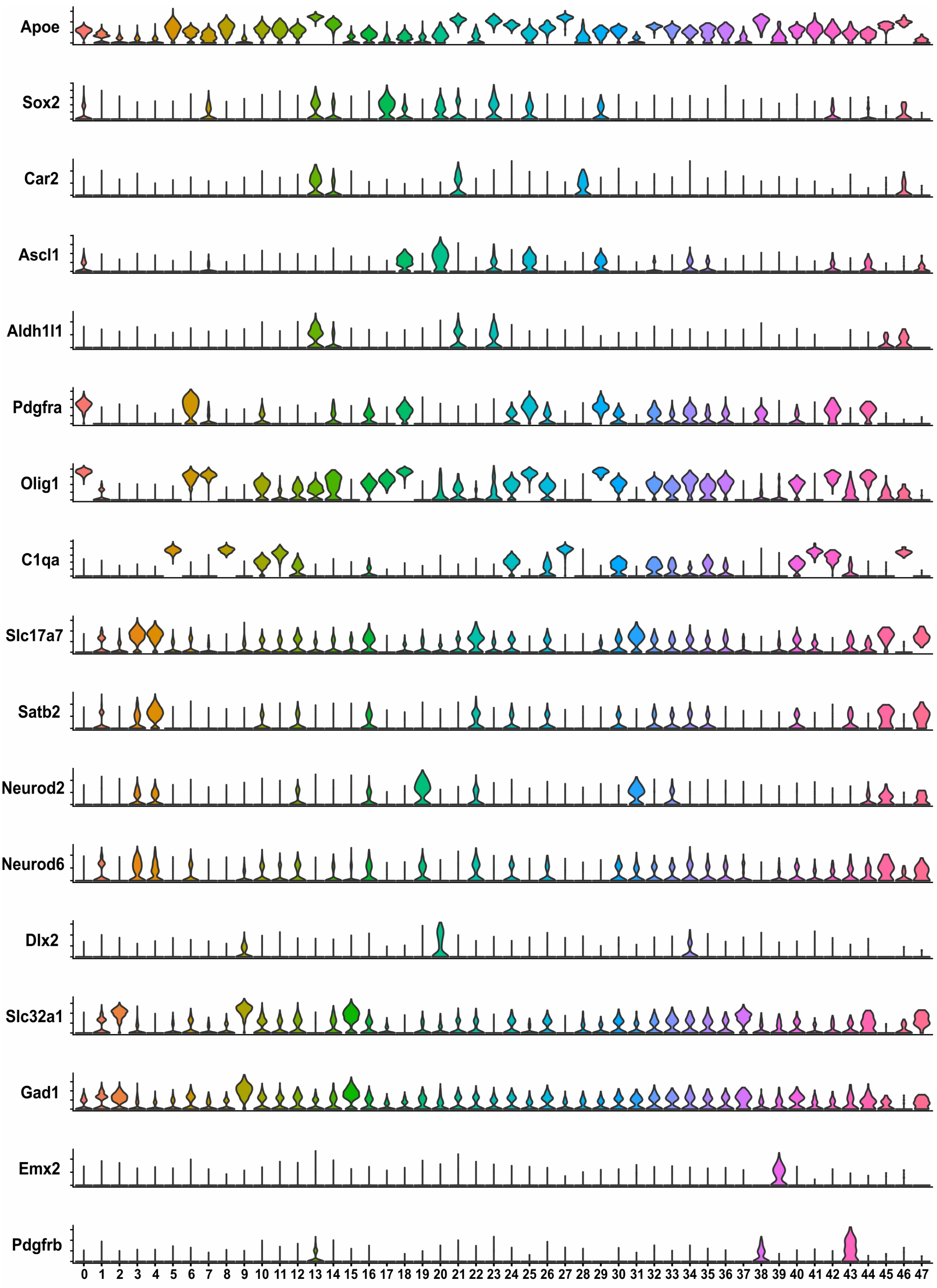


Zhu *et al.*, Figure 1-Figure supplement 3

**Figure 1-S3: Neuron selection**

Violin plots illustrating specific gene expression in various clusters. Cluster 1, 2, 3, 4, 9, 12, 15, 16, 19, 22, 31, 37, 45 and 47 were categorized as neurons.

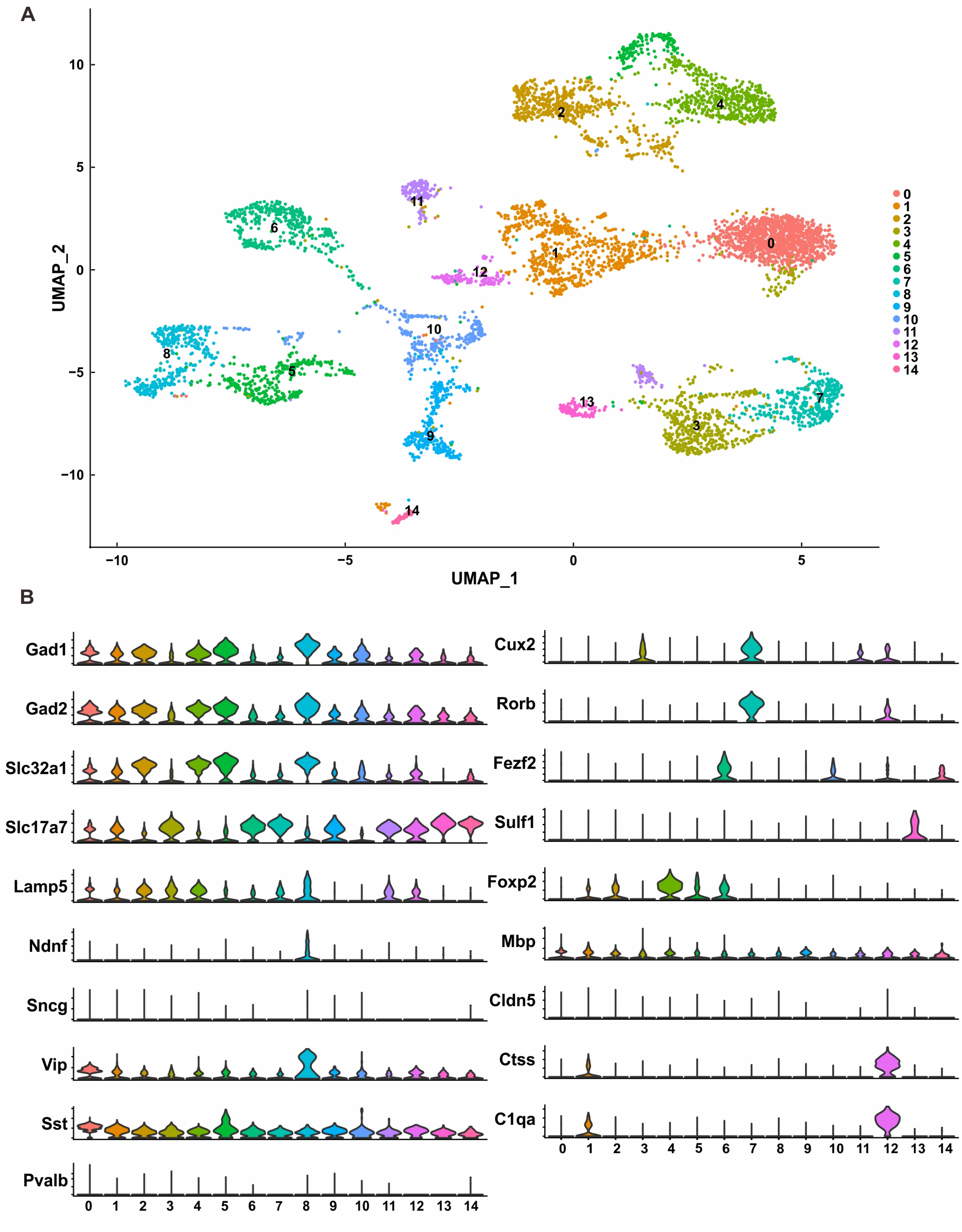


**Zhu *et al.*, Figure 1-Figure supplement 4**

**Figure 1-S4: Neuron clusters**

(**A**) UMAP analysis of neurons (n=6505). (**B**) Violin plot displaying specific gene expression in various neuron clusters. Cluster 0, 1, 2, 4, 5, 8, and 10 were identified as inhibitory neurons; cluster

3, 7, and 11 were categorized as upper layer excitatory neurons; cluster 6, 13, 14 were classified as deep layer excitatory neurons; cluster 9 was labeled as unidentified excitatory neurons; and cluster 12 did not exhibit typical neuron characteristics.

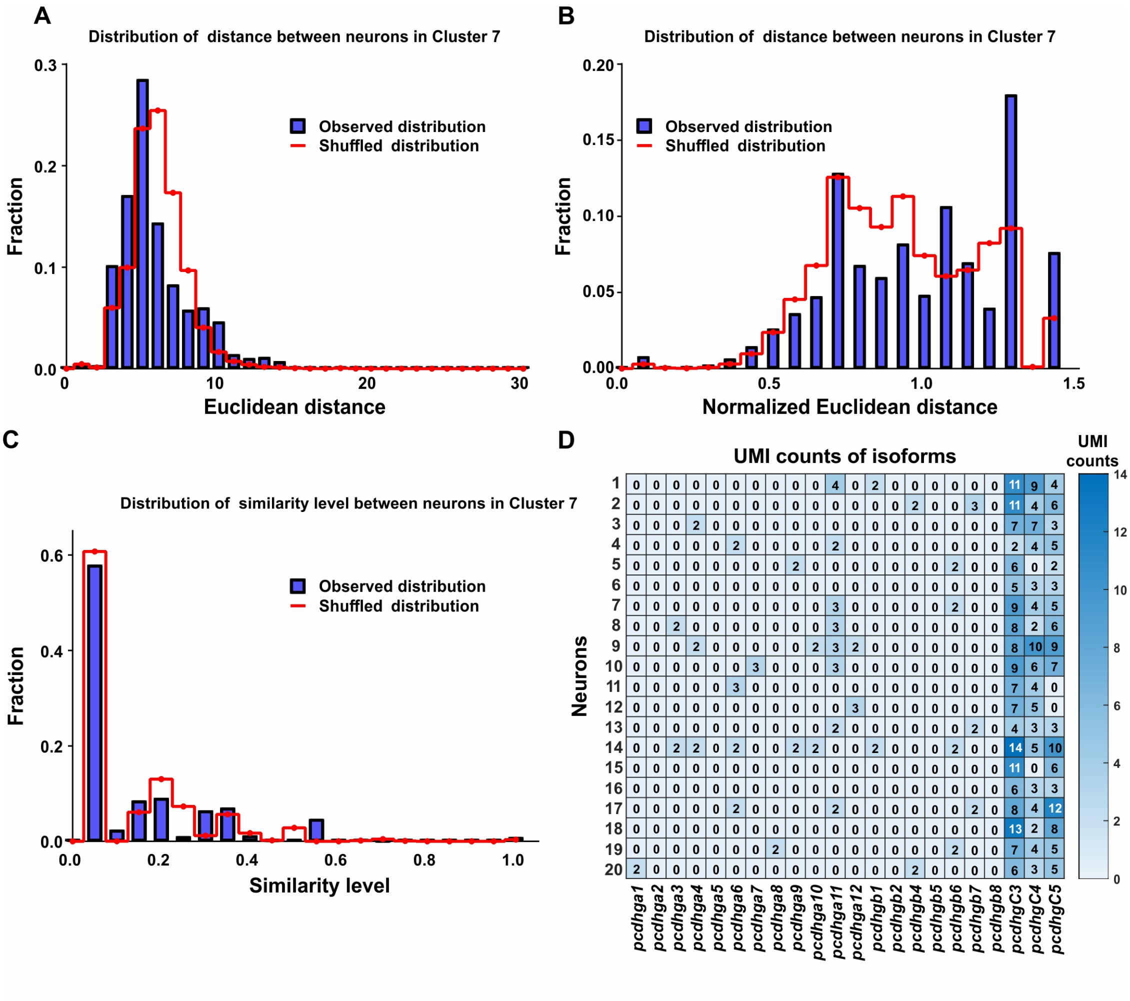


Zhu *et al.*, Figure 1-Figure supplement 5

**Figure 1-S5: Diversely combinatorial expression of γ-PCDH in L2/3 neurons of Cluster 7**

(**A**) Observed distribution (blue) of Euclidean distance between neurons from Cluster 7 in comparison to the distance from shuffled data (red). (**B**) Observed distribution (blue) of normalized Euclidean distance between neurons from Cluster 7 in comparison to the distance from shuffled data (red). (**C**) The similarity level between cells in Cluster 7 under both observed (blue) and shuffled (red) conditions. (**D**) The detailed expression for randomly selected 20 neurons in Cluster 7. Each row represents one neuron, and each column denotes one isoform. The numbers in the box, as well as the colors, show the RNA counts measured by UMI.

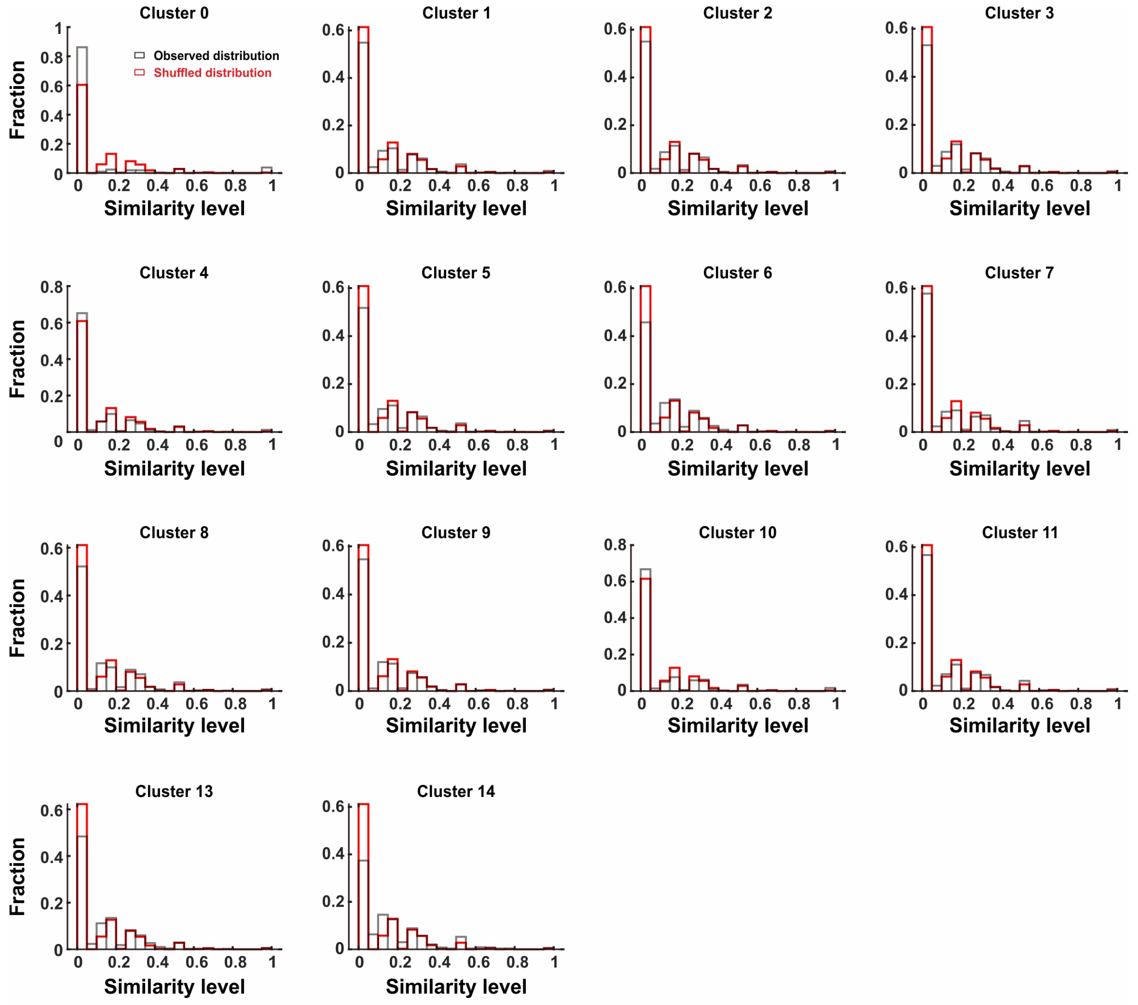


Zhu *et al.*, Figure 1-Figure supplement 6

**Figure 1-S6: Distribution of similarity levels among cell pairs across different clusters.** Gray and red bars indicate observed and shuffled data. The panel “Cluster 7” showed the same dataset with Fig.1-S5C for the consistency.

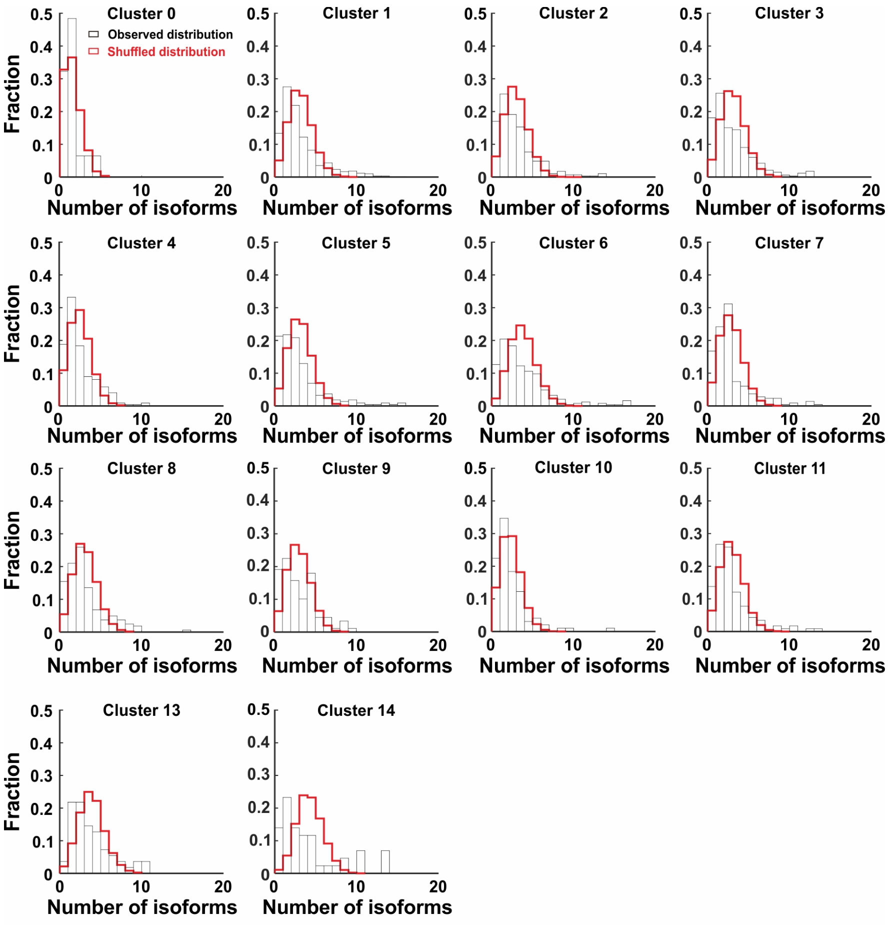


Zhu *et al.*, Figure 1-Figure supplement 7

**Figure 1-S7: A weak but significant co-occurrence of γ-PCDH variable isoforms in most of neocortical neurons (except neurons in cluster 0).**

Fraction distribution of the number of the expressed isoforms per cell in different clusters.

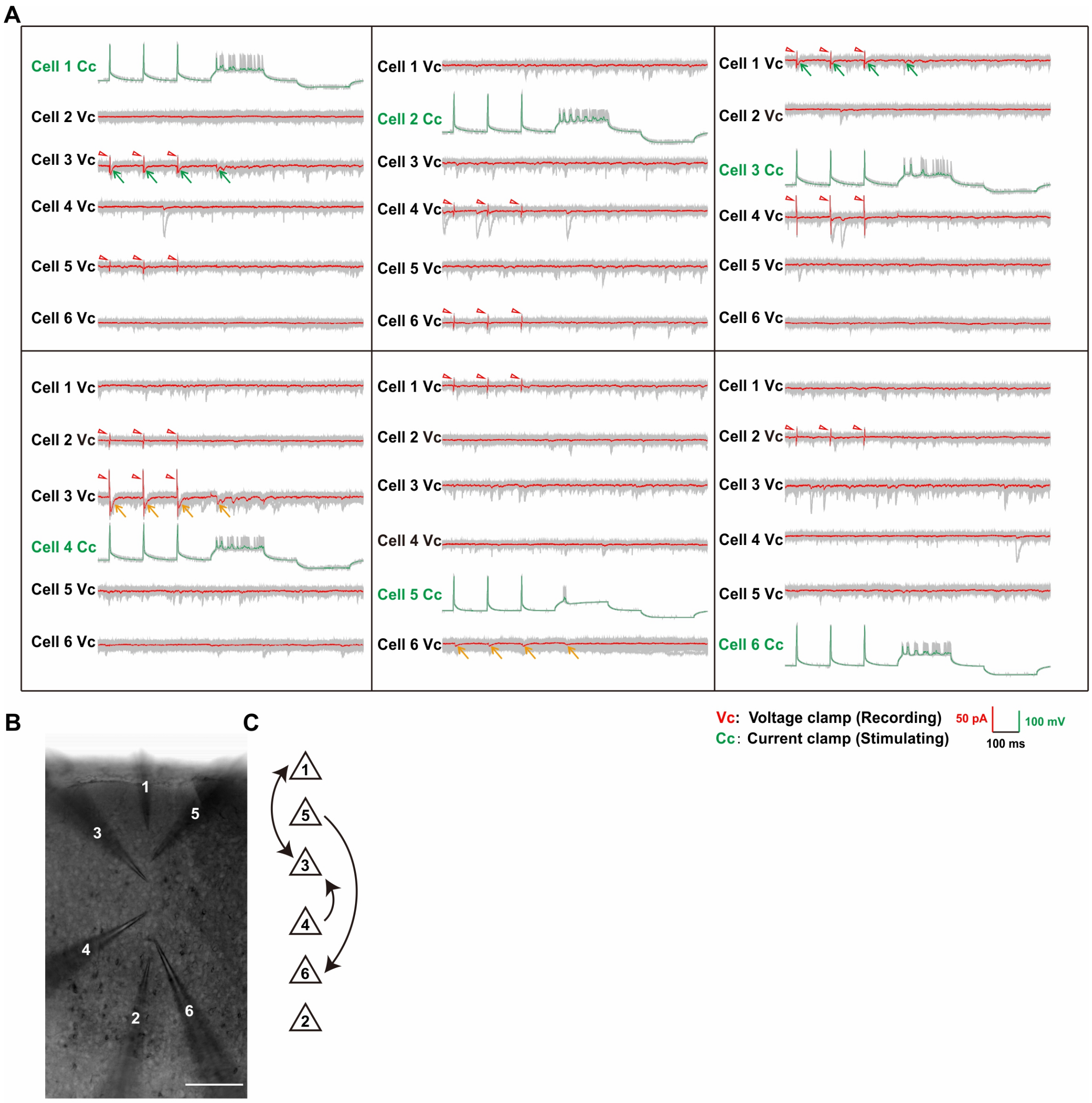


Zhu *et al.*, Figure 2-supplement 1

**Figure 2-S1: Multi-electrode whole-cell patch-clamp recording.**

(**A**) Full sample traces of electrophysiological recording showed in Fig2A. Individual traces are shown in gray and the averages from 10 trials are shown in red. Positive evoked postsynaptic responses are indicated by arrows. Orange/green arrows: unidirectional/ bidirectional connections. Stimulus artifacts are pointed out by red triangles. VC: cell was recorded under the voltage clamp; CC: cell was recorded under the current clamp. (**B**) Images taken by the differential interference microscope for the recorded 6 neurons and their corresponding electrodes in (**A**). (**C**) Schematic of the synaptic connectivity among these 6 neurons showed in (**A**) and (**B**).

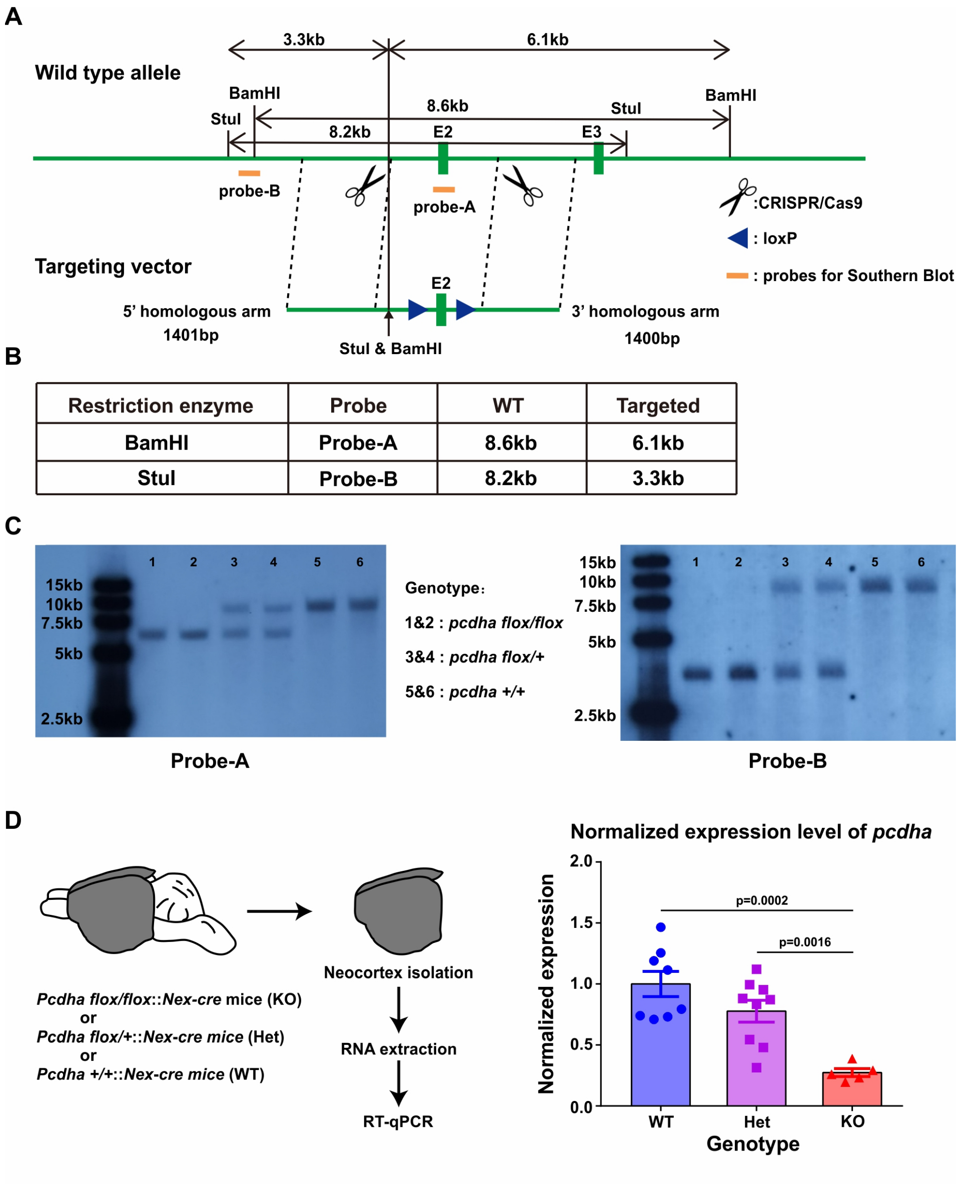


Zhu *et al.*, Figure 2-Figure supplement 2

**Figure 2-S2: Generation and characterization of *Pcdha*$^{flox/flox}$ mice.**

(**A**) Strategy of introducing loxP sites at both sides of the second exon of *Pcdha* using CRISPR/Cas9. The orange bars indicate the sites for the Southern Blot assay. The left cutting site is caused by sgRNA_1, the right site is caused by sgRNA_2. (**B**) The theoretical lengths in the Southern Blot assay for both probes. (**C**) Results of Southern Blot for both probe A (left) and probe B (right) from six mice. (**D**) The reduction of *Pcdha* expression in *Pcdha* CKO mice as revealed by qRT-PCR. Left: experiment design. The entire neocortex was dissected for RNA extraction. Notably, the neocortex was isolated from mice where the Nex-cre gene was expressed specifically in excitatory neurons within the neocortex. Right: qRT-PCR quantification of *Pcdha* expression in *Pcdha* KO mice cortex.

Student's *t* test was used for statistical analysis.

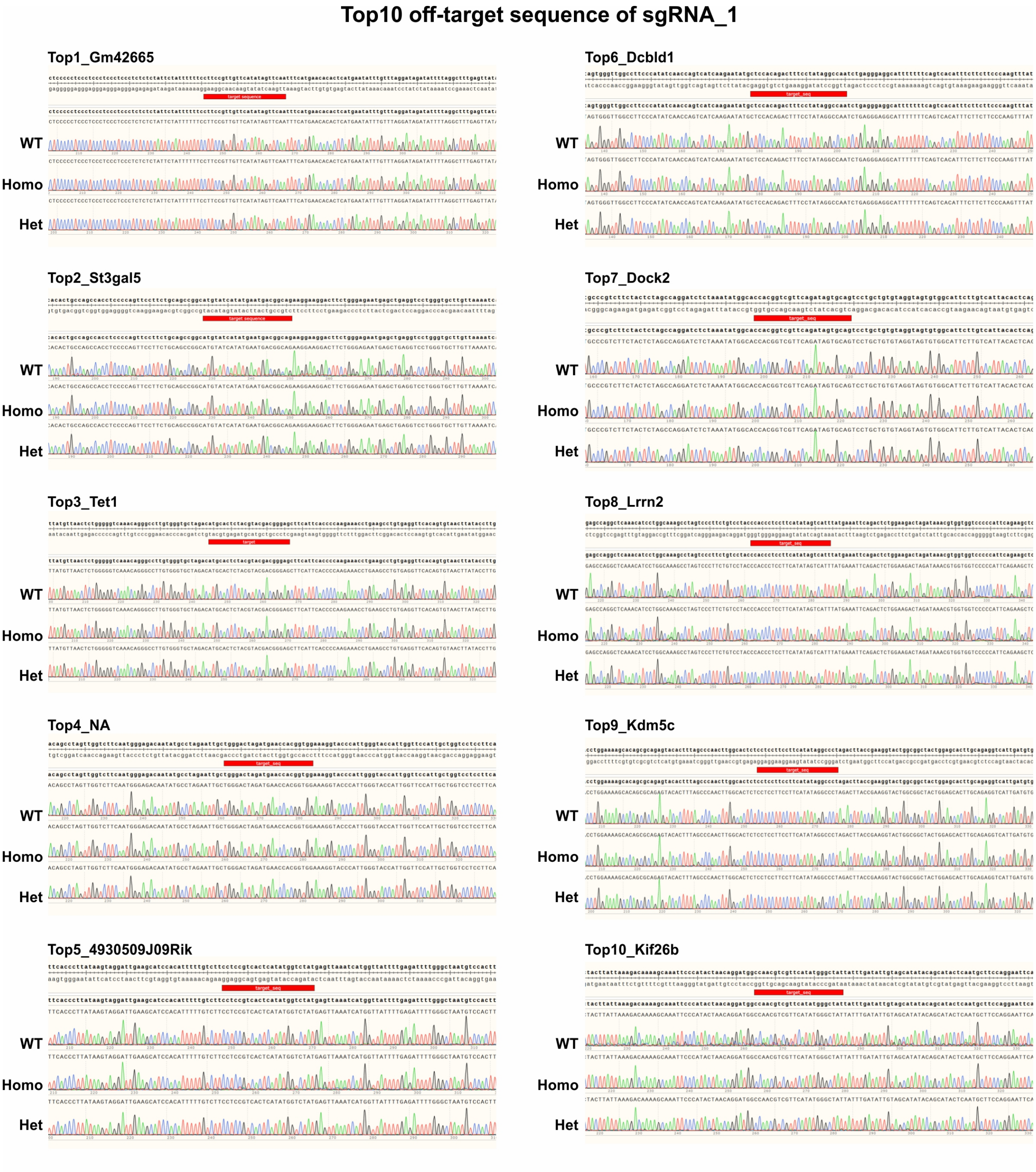


Zhu *et al.*, Figure 2-Figure supplement 3

**Figure 2-S3: Top 10 off-target sequences of sgRNA_1.**

Sanger sequencing results of top 10 off-target regions. WT: *pcdha*$^{+/+}$ mice. Homo: *pcdha*$^{flox/flox}$ mice. Het: *pcdha*$^{+/flox}$ mice. The mice used were littermates. The red bars (target_seq) showed the predicted off-target regions in the mouse genome.

**Top10 off-target sequence of sgRNA_2**

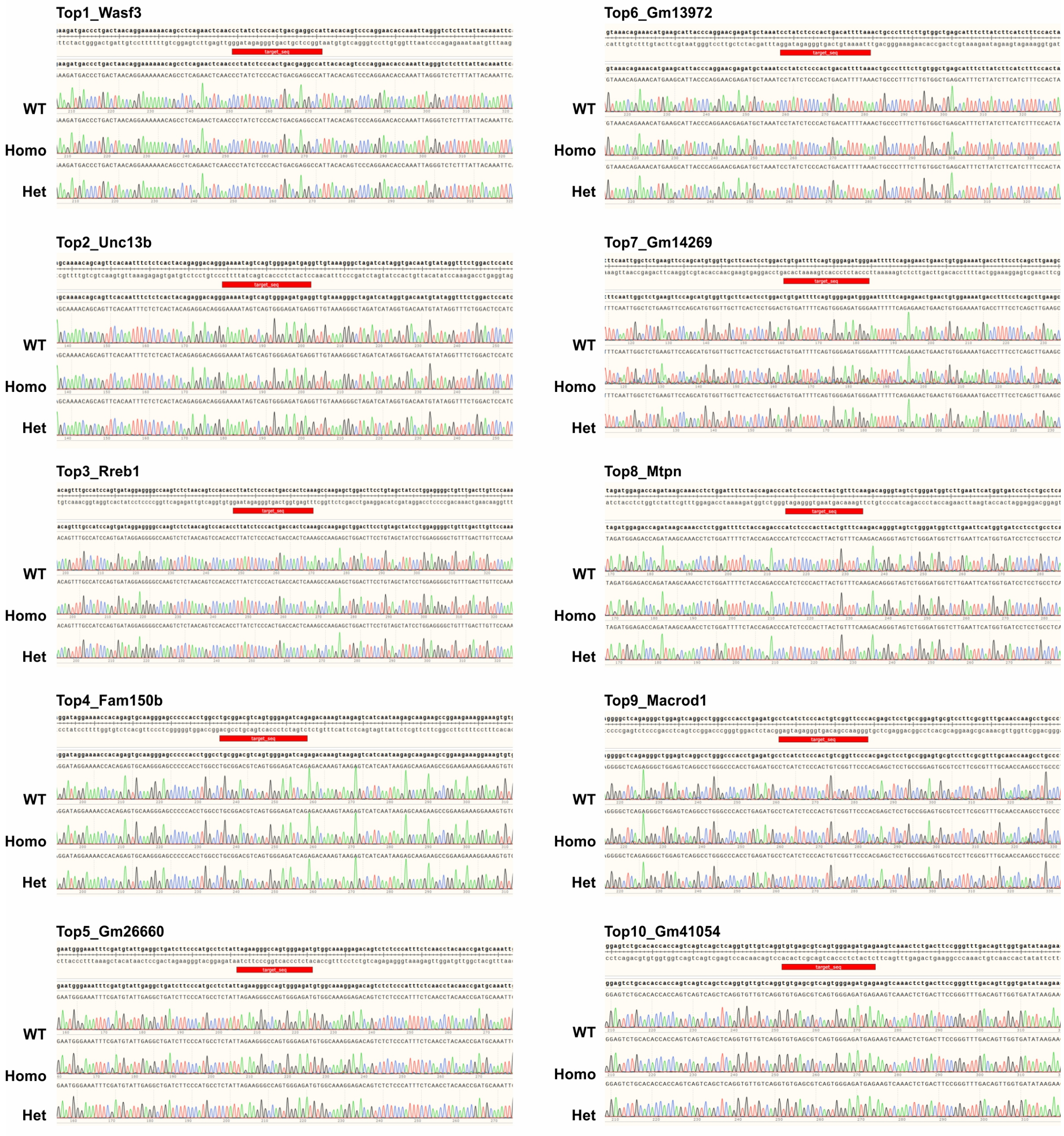


Zhu *et al.*, Figure 2-Figure supplement 4

**Figure 2-S4: Top 10 off-target sequences of sgRNA_2.**

Sanger sequencing results of top 10 off-target regions. WT: *pcdha*$^{+/+}$ mice. Homo: *pcdha*$^{flox/flox}$ mice. Het: *pcdha*$^{+/flox}$ mice. The mice used were littermates. The red bars (target_seq) showed the predicted off-target regions in the mouse genome.

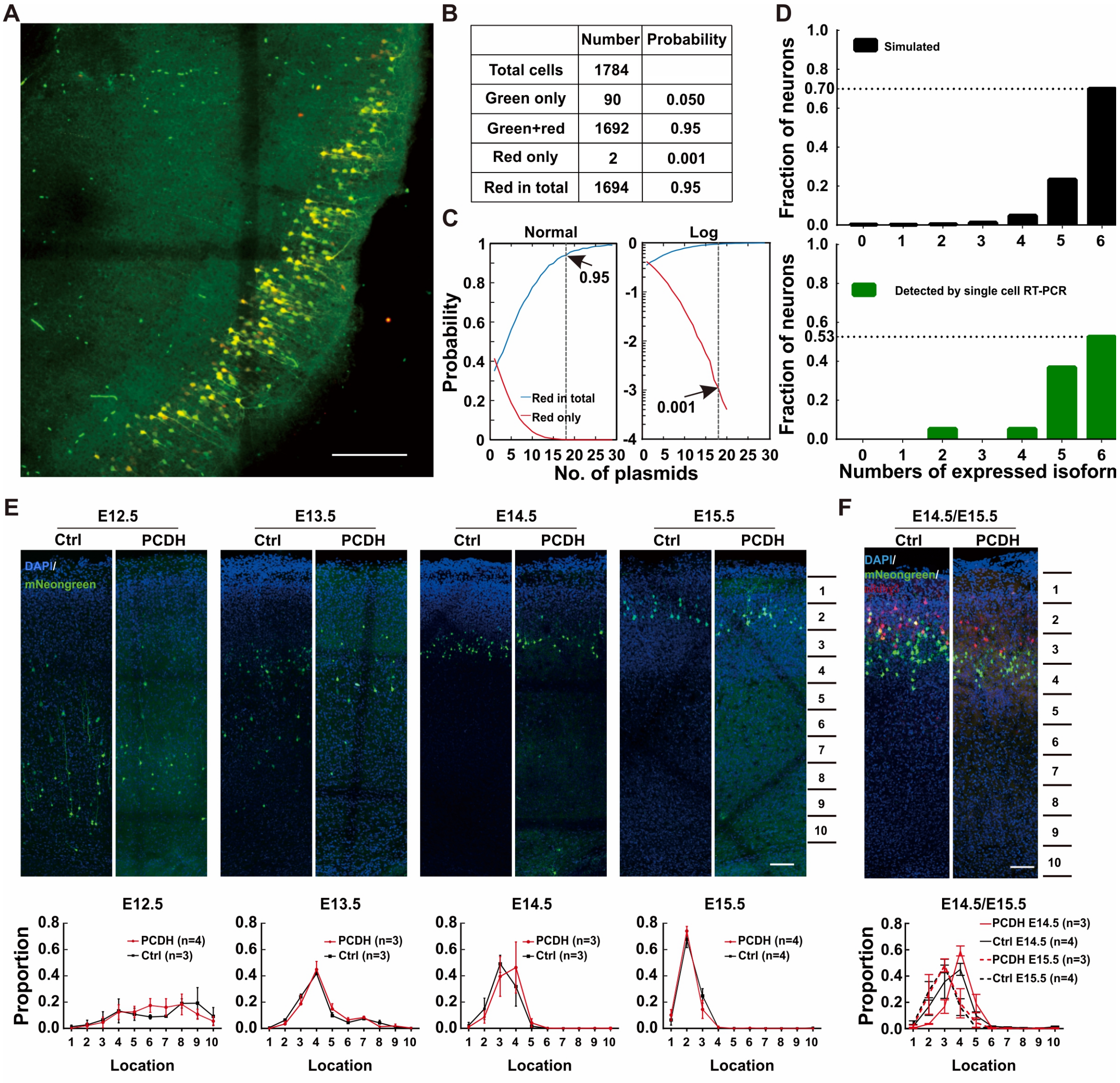


**Zhu *et al.*, Figure 3-Figure supplement 1**

**Figure 3-S1: Characterizing the overexpression effect after electroporation.**

(**A**) A sample confocal image of electroporated neurons with five different isoforms tagged with mNeongreen and the sixth with mCherry. Scale bar =100µm. (**B**) The numbers of cells and corresponding probabilities in total population after an electroporation containing five different isoforms with green and the sixth with mCherry. (**C**) The normal (top) and logarithmic (bottom) probability distribution of plasmid numbers in electroporated neurons after the simulation of 10,000 cells. The top panel shows the correlation between the plasmid number and the probability of "red in total" (blue line), while the bottom panel shows the correlation between the plasmid number and the likelihood of "red only" (red line). (**D**) Distribution of the expressed isoform numbers from the stimulation (upper panel) or from the single-cell RT-PCR (lower panel). (**E**) Overexpressing γ-PCDH in neurons did not affect their layer distributions in the neocortex. Sample confocal images for the distribution of neurons overexpressing γ-PCDH A2 through electroporation at different embryonic stages. The lower panels demonstrate the numbers of neurons in different evenly distributed zones of the neocortex along the pial surface to the edge of white matter. Scale bar: 100 µm. (**F**) The distribution of neurons after a sequential-electroporation with four plasmids γ-PCDH A4, A9, A11, or B6 tagged by mNeongreen at E14.5 and other four γ-PCDH A2, A6, A8, or B1 tagged by mRuby3 at E15.5. The lower panel shows the distribution of cell numbers in different zones (Zone 1 starts on the top of the pial surface and zone 10 ends at the border with the white matter). Scale bar: 100 µm. Two-way ANOVA was used for statistical analysis in **E** and **F**.

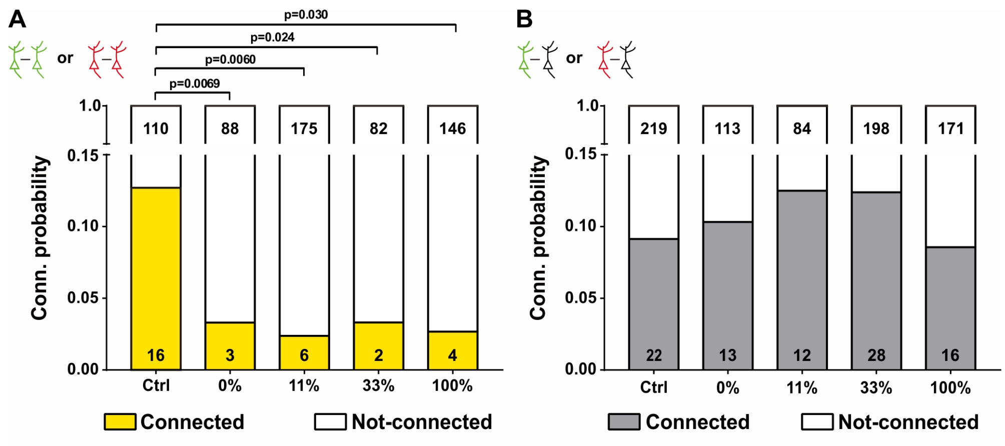


Zhu *et al.*, Figure 4-Figure supplement 1

**Figure 4-S1: The effect of overexpressing γ-PCDHs on synaptic connectivity.**

(**A**) The connectivity probability for neuron pairs overexpressed same γ-PCDHs combinations (labeled with the same fluorescence). The statistical difference was compared between each group and the control group. (**B**) The connectivity probability for neuron pairs contained one overexpressed cell and one adjacent cell without fluorescence. Chi-square test and false discovery rate (FDR) correction were used to determine the statistical difference in (**A**) and (**B**).
